# Network analysis for identifying potential anti-virulence targets through whole transcriptome analysis of *Pseudomonas aeruginosa* and *Staphylococcus aureus* exposed to certain anti-pathogenic polyherbal formulations

**DOI:** 10.1101/2023.04.27.538520

**Authors:** Feny Ruparel, Siddhi Shah, Jhanvi Patel, Nidhi Thakkar, Gemini Gajera, Vijay Kothari

## Abstract

Transcriptome of two important pathogens, *Pseudomonas aeruginosa* and *Staphylococcus aureus* exposed to two different quorum-modulatory polyherbal formulations were subjected to network analysis to identify the most highly networked differentially expressed genes (hubs) as potential anti-virulence targets. Genes associated with denitrification and sulfur metabolism emerged as the most important targets in *P. aeruginosa*. Increased build-up of nitrite (NO2) in *P. aeruginosa* culture exposed to the polyherbal formulation *Panchvalkal* was confirmed through *in vitro* assay too. Generation of nitrosative stress and inducing sulfur starvation seems to be effective anti-pathogenic strategies against this notorious gram-negative pathogen. Important targets identified in *S. aureus* were the transcriptional regulator *sarA*, immunoglobulin-binding protein Sbi, serine protease *SplA*, the *saeR/S* response regulator system, and gamma-haemolysin components *hlgB* and *hlgC*. Further validation of the potential targets identified in these pathogens is warranted through appropriate *in vitro* and *in vivo* assays in model hosts. Such validated targets can prove vital to many antibacterial drug discovery programmes globally.

## Introduction

Despite wide recognition of AMR (antimicrobial resistance) as a major global health threat, the progress on front of discovery and development of new antibiotics in last 3-4 decades clearly has fallen short from being satisfactory. For a variety of reasons e.g., lack of interest among major pharmaceutical firms, rapid emergence and spread of resistance among pathogenic bacterial populations, dearth of new validated cellular and molecular targets, the list of effective antimicrobials available for treatment of resistant infections remains short. The status of antibiotic discovery research has been reviewed thoroughly by Årdal et al. (2018), Årdal et al. (2020), Prasad et al. (2022), and Årdal et al. (2023). Since most currently available antibiotics target a narrow range of bacterial traits i.e., cell envelope synthesis, protein or nucleic acid synthesis, or folic acid synthesis, a truly new class of antibiotic will be discovered only if we have a longer list of validated targets. Development of new bactericidal antibiotics is not the only way of tackling the slow pandemic of AMR infections; discovery of resistance modifiers and non-antibiotic virulence-attenuating agents can also be of great value [Laxminarayan et al., (2013); Cheng et al., (2016)]. Hence identification of new potential targets for bactericidal antibiotics as well as antibiotic adjuvants both is useful. There is a clear desperate need for antibiotics with previously unexploited new targets and wide target diversity in the discovery pipeline. One of the major challenges in way of antibacterial discovery is associated with the proper target selection, e.g., the requirement of pursuing molecular targets that are not prone to rapid resistance development [Silver (2011)].

Various public health agencies like CDC (Centers for Disease Control and Prevention, USA), WHO (World Health Organization), DBT (Department of Biotechnology, India) etc. have published lists of priority pathogens against whom novel antimicrobials need to be discovered urgently. Antibiotic-resistant strains of *Pseudomonas aeruginosa* and *Staphylococcus aureus* commonly appear on all such lists. As per CDC’s Vital Signs report (https://www.cdc.gov/vitalsigns/index.html) more than 33% of the bloodstream infections in patients on dialysis in the US in 2020 were caused by *S. aureus*. This gram-positive human commensal has been recognised as an important opportunistic pathogen responsible for wide range of infections [Kenny et al. (2009)]. *P. aeruginosa* is the primary cause of gram-negative nosocomial infections. Its ability to adapt to a wide range of environmental niches combined with its nutritional versatility and genome plasticity, along with a multitude of intrinsic and acquired resistance mechanisms make it one of the most notorious pathogen of critical clinical importance. Efforts for finding perturbants capable of targeting the *P. aeruginosa* pathogenicity and antibiotic resistance are highly desired [Langendonk et al., (2021)].

We had previously studied the anti-virulence effect of certain polyherbal formulations against *S. aureus* or *P. aeruginosa* at the whole transcriptome level of the target pathogen, wherein we gained some insight into the molecular mechanisms associated with the virulence-attenuating potential of the test formulations, which was largely independent of any growth-inhibitory effect. Pathogens exposed to the test formulations were compromised in their ability to kill the model host *Caenorhabditis elegans*. The current study attempted network analysis of the differentially expressed genes of *P. aeruginosa* and *S. aureus* exposed to the anti-pathogenic polyherbal formulations *Panchvalkal* (Joshi et al., 2019) and Herboheal (Patel et al., 2019) respectively, reported in the previous studies, with an aim to identify highly networked genes as potential anti-virulence targets.

## Methods

### Network analysis

We accessed the list of differentially expressed genes (DEG) for *Panchvalkal* (PentaphyteP-5^®^)-exposed *P. aeruginosa* (NCBI Bioproject ID 386078) and Herboheal -exposed-*S. aureus* (NCBI Bioproject ID 427073). The *P. aeruginosa* used was a multidrug-resistant strain. Network analysis for both the studies was carried out independently, wherein only the DEG fulfilling the dual filter criteria of log fold-change ≥2 and FDR≤0.01 were selected for further analysis. The list of such DEG was fed into the database STRING (v.11.5) [Szklarczyk et al., (2019)] for generating the PPI (Protein-Protein Interaction) network. Then the genes were arranged in decreasing order of ‘node degree’ (a measure of connectivity with other genes or proteins), and those above a certain threshold value were subjected to ranking by cytoHubba (v.3.9.1) [Chin et al., (2014)]. Since cytoHubba uses 12 different ranking methods, we considered the DEG being top-ranked by ≥6 different methods (i.e., 50% of the total ranking methods) for further analysis. These top-ranked shortlisted proteins were further subjected to network cluster analysis through STRING and those which were part of multiple clusters were considered ‘hubs’ which can be taken up for further validation of their targetability. Here ‘hub’ refers to a gene or protein interacting with many other genes/ proteins. Hubs thus identified were further subjected to co-occurrence analysis to see whether an anti-virulence agent targeting them is likely to satisfy the criterion of selective toxicity (i.e., targeting the pathogen without harming host). This sequence of analysis allowed us to end with a limited number of proteins which satisfied various statistical and biological significance criteria simultaneously i.e. (i) log fold-change ≥2; (ii) FDR≤0.01; (iii) relatively higher node degree; (iv) top-ranking by at least 6 cytoHubba methods; (v) (preferably) member of more than 1 local network cluster; and (vi) high probability of the target being absent from the host. A schematic presentation of the methodology employed for network analysis is presented in Figure 1.

**Figure 1:**
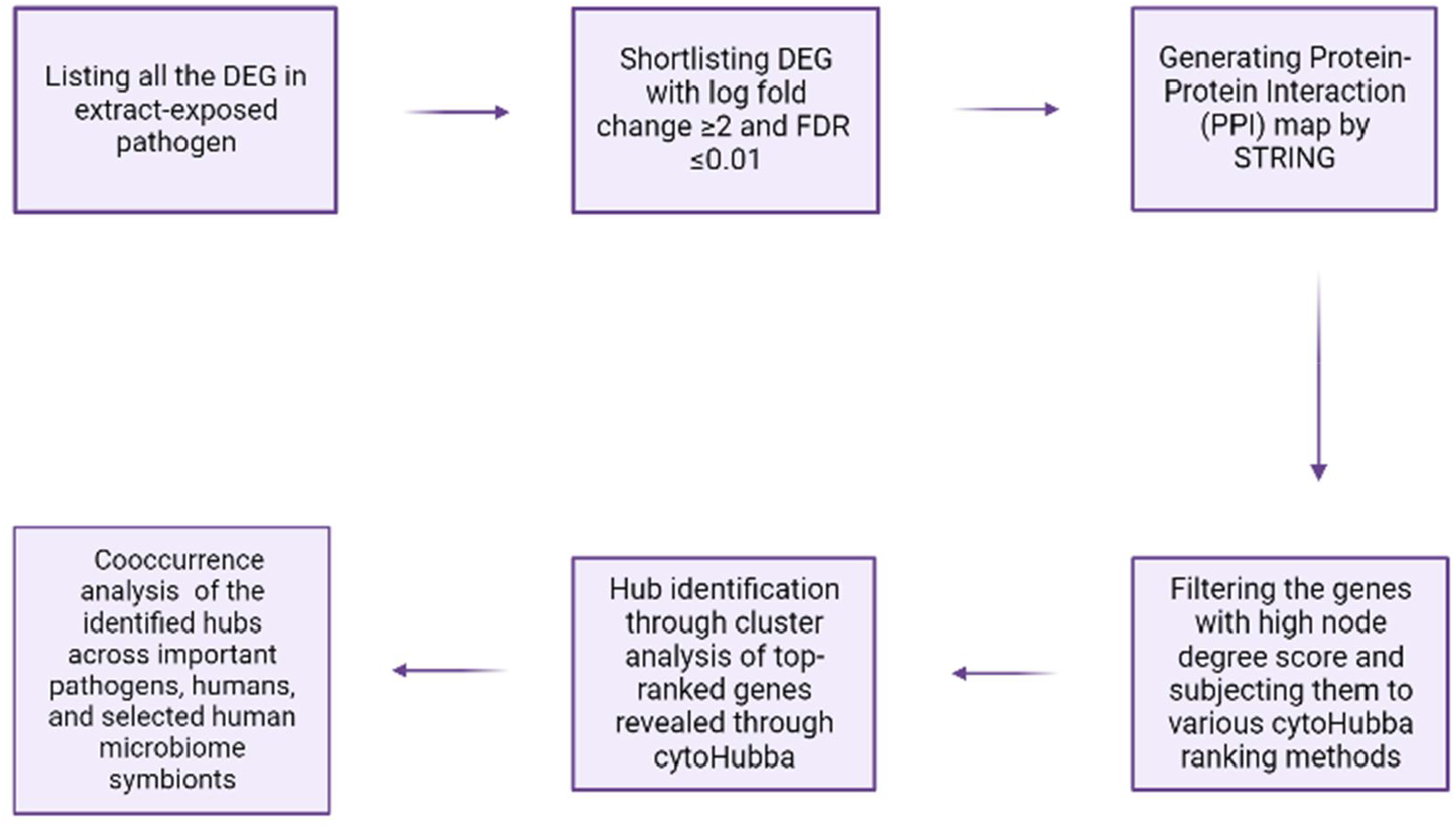
A schematic of methodology for network analysis and hub identification. (Image created through BioRender: https://app.biorender.com/)

### Nitrite estimation

Nitrite estimation in *P. aeruginosa* culture supernatant was done through Griess assay (Misko et al., 1993). *P. aeruginosa* was grown in Pseudomonas Broth (HiMedia, Mumbai) with or without *Panchvalkal* (567µg/mL; dried extract powder without any bulking agent was procured from Dr. Palep’s Medical Education and Research Foundation Pvt. Ltd., Mumbai, India, and dissolved in DMSO for assay purpose) at 35°C for 21±1 h. Following incubation, cell density was quantified at 764 nm (Joshi et al., 2016), and then the bacterial culture suspension was centrifuged at 13,600 g for 10 min. Resulting supernatant was mixed with Griess reagent (Sigma-Aldrich) in 1:1 ratio, and incubated for 15 min in dark at room temperature. Absorbance of the resulting pink colour was quantified at 540 nm (Agilent Technology Cary 60 UV-Vis). These OD values were plotted on standard curve prepared using NaNO2 to calculate the nitrite concentration. To nullify any effect of variation in cell density between control and experimental culture, Nitrite Unit (i.e., nitrite produced per unit of growth) was calculated by dividing the nitrite concentration values by cell density. Sodium nitroprusside (Astron chemicals, Ahmedabad) being a chemical known to be capable of generating nitrosative stress in bacteria (Barraud et al., 2006; Barnes et al., 2013; Fida et al., 2018) was used as a positive control. Appropriate vehicle control [i.e., bacteria grown in presence of 0.5%v/v DMSO (Merck)], negative control (deionized water), and abiotic control (*Panchvalkal* supplemented Pseudomonas broth) were included in the experiment. Griess reagent was added in all these controls in the same proportion as that in extract-exposed or not-exposed bacterial culture samples.

## Results

### Network analysis of DEG in *Panchvalkal-*exposed *P. aeruginosa*

Our original experimental study exposed *P. aeruginosa* to *Panchvalkal* at 567 µg/mL, wherein the extract-exposed pathogen could kill 90% lesser host worms than its extract-not-exposed counterpart. Whole transcriptome study revealed that approximately 14% of the *P. aeruginosa* genome was expressed differently under the influence of *Panchvalkal*. The total number of DEG satisfying the dual criteria of log fold-change ≥2 and FDR≤0.01 was 228, of which 105 were downregulated (Table S1), and 123 were up-regulated (Table S4). We created PPI network for up- and down-regulated genes separately (Figure 5 and Figure 2 respectively). PPI network for downregulated genes generated through STRING is presented in Figure-2, which shows 101 nodes connected (105 genes were fed to string, out of which 101 were shown in the PPI network) through 86 edges with an average node degree of 1.7. Since the number of edges (86) in this PPI network is 3.18-fold higher than expected (27) with a PPI enrichment p-value <1.0e-16, this network can be said to possess significantly more interactions among the member proteins than what can be expected for a random set of proteins of the identical sample size and degree distribution. Such an enrichment can be taken as an indication of the member proteins being at least partially biologically connected. When we arranged the 105 downregulated genes in decreasing order of node degree, 52 nodes were found to have a non-zero score (Table S2), and we selected top 13 genes with a node degree ≥6 for further ranking by different cytoHubba methods. Then we looked for genes which appeared among top-10 ranked candidates by ≥6 cytoHubba methods, and 10 such shortlisted genes (Table S3) were further checked for interactions among themselves followed by cluster analysis (Figure-3), which showed them to be strongly networked as the average node degree score was 8. This network possessed 40 edges as against expected (zero) for any such random set of proteins (PPI enrichment p-value <1.0e-16). The PPI network generated through STRING showed these 10 important genes to be distributed among three different local network clusters. Five (norB, norC, norD, nirS, and nirQ) of the predicted hubs were part of each of the three clusters, and they have role in denitrification (Arai, 2011). Of the remaining 5 predicted hub proteins, one more (NorE) is also associated with nitrogen metabolism, and two (nosL and nosY) have role in denitrification as well as copper homeostasis. These three proteins were members of two out of three clusters. The eight proteins (Table 1) found to be members of minimum two clusters can be said to be potential hubs, whose down-regulation can be hypothesized to attenuate *P. aeruginosa* virulence.

**Table 1.** Hubs identified as potential targets from among the down-regulated genes in *Panchvalkal*-exposed *P. aeruginosa*.

| No. | Gene ID | Gene Name | Functional Role |
| --- | --- | --- | --- |
| 1 | PA0520 | *nirQ* | Denitrification regulatory protein NirQ |
| 2 | PA0519 | *nirS* | Heme d1 biosynthesis protein, which is important for denitrification (Adamczack et al., 2014) |
| 3 | PA0524 | *norB* | Nitric oxide reductase subunit B |
| 4 | PA0523 | *norC* | Nitric oxide reductase subunit C |
| 5 | PA0525 | *NorD* | Nitric oxide reductase NorD protein |
| 6 | PA0521 | *NorE* | Nitric oxide reductase NorE protein |
| 7 | PA3395 | *nosY* | Nitrous oxide reductase; a Cu-processing system permease protein having role in denitrification pathway (Dettman et al., 2013) |
| 8 | PA3396 | *nosL* | A lipoprotein attached to the outer membrane described as a copper-binding protein. Regulator of *nos* operon, NosR also associates with NosL. This protein is probably responsible for the insertion and coordination of the multicopper centre within NosZ (Borrero-de Acuña et al., 2017). |

**Figure 2.**
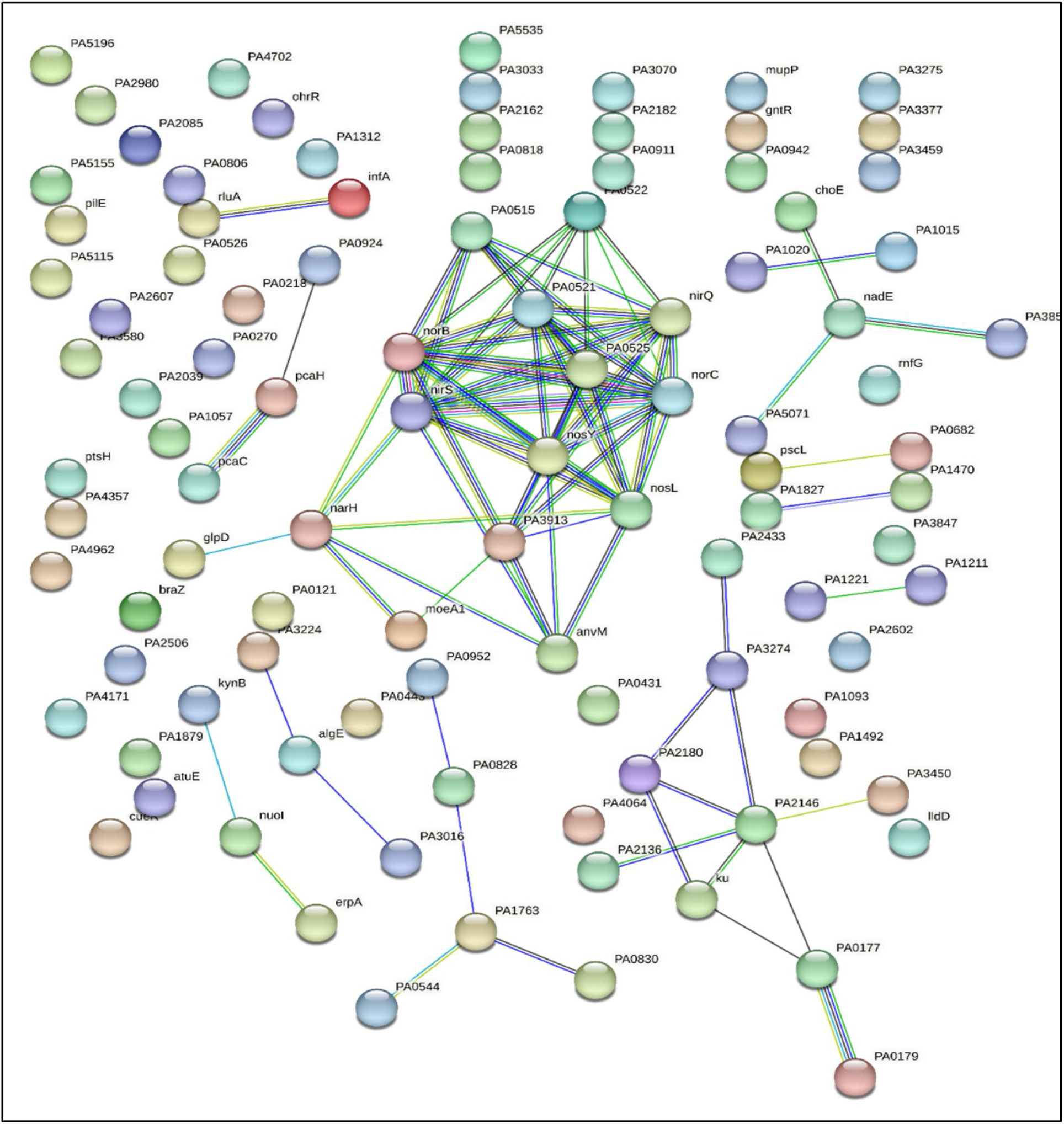
Protein-Protein Interaction (PPI) network of down regulated genes in *Panchvalkal* exposed *P. aeruginosa*. Edges represent protein-protein associations that are meant to be specific and meaningful, i.e. proteins jointly contribute to a shared function; this does not necessarily mean they are physically binding to each other.Network nodes represent proteins. Splice isoforms or post-translational modifications are collapsed, i.e. each node represents all the proteins produced by a single, protein-coding gene locus.

**Figure 3.**
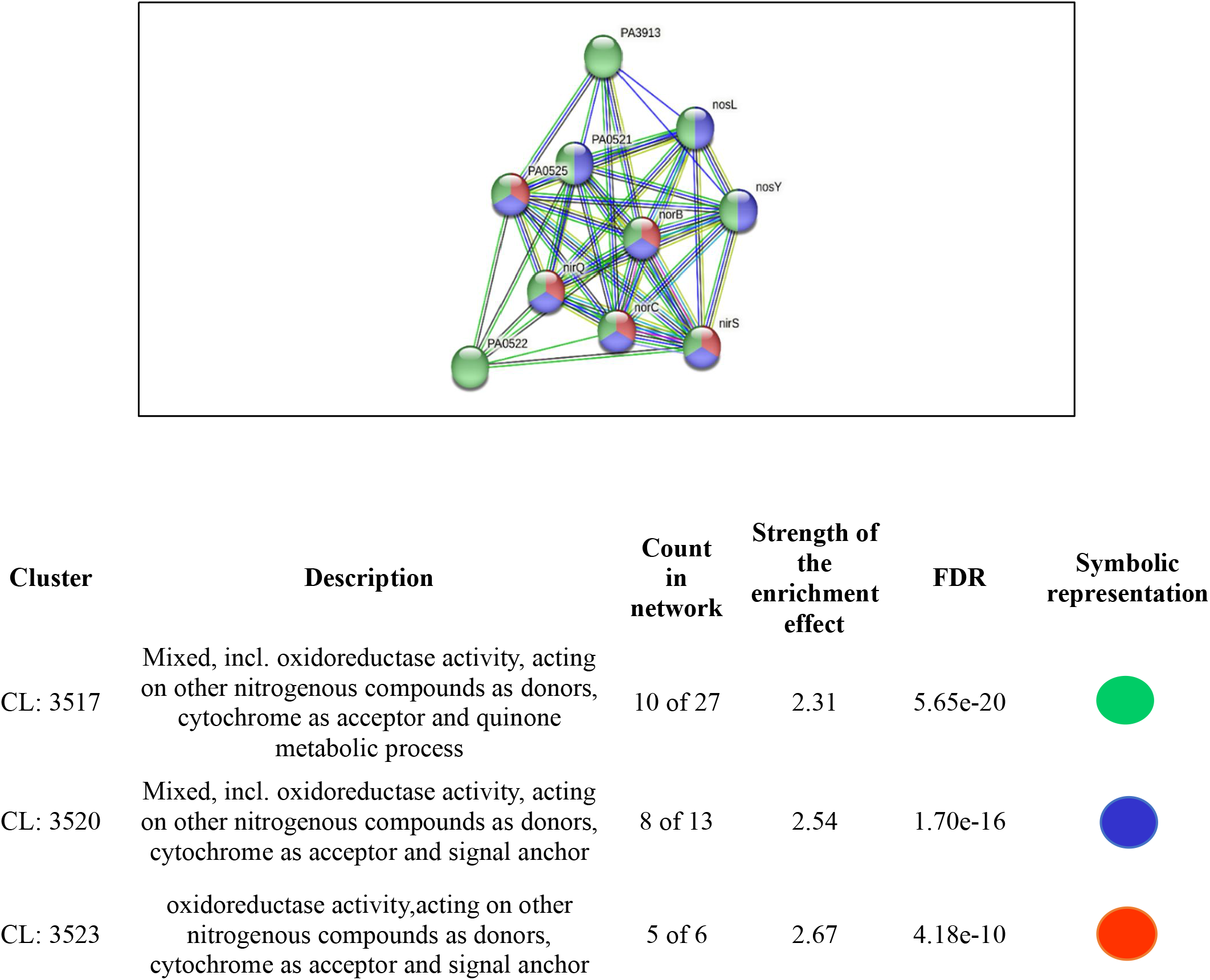
PPI network of top-ranked genes revealed through cytoHubba among down-regulated DEG in *Panchvalkal*-exposed *P. aeruginosa*.

Since all the targets mentioned in Table-1 are known to play important role in *P. aeruginosa* with respect to detoxification of reactive nitrogen species, we hypothesized that *Panchvalkal*-treated *P. aeruginosa’*s ability to detoxify reactive nitrogen species is compromised. To check this hypothesis, we quantified nitrite concentration in extract-treated *P. aeruginosa* culture, wherein it was found to have 31% higher nitrite concentration in supernatant as compared to control (Figure 4). This higher accumulation of nitrite can be taken as an indication of compromised denitrification efficiency as nitrite is an intermediate of denitrification pathway (Borrero et al., 2017).

**Figure 4.**
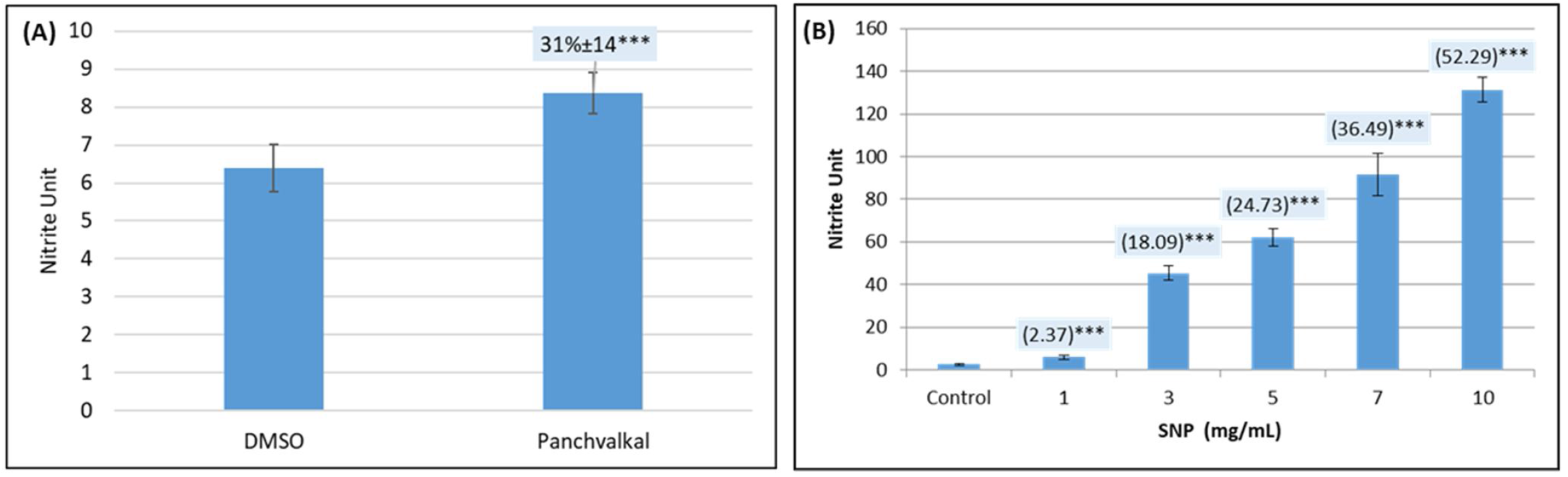
*Panchvalkal*-treated *P. aeruginosa* culture has higher extracellular accumulation of nitrite. While nitrite concentration in vehicle control (*P. aeruginosa* incubated in media supplemented with 0.5%v/v DMSO) was at par to that without DMSO, *Panchvalkal* caused nitrite concentration in *P. aeruginosa* culture supernatant to rise (Fig. 4A). Sodium nitroprusside used as positive control caused a dose-dependent 2.37 to 52.29-fold higher nitrite build up in *P. aeruginosa* culture (Fig. 4B). Nitrite Unit (i.e., Nitrite concentration: Cell density ratio) was calculated to nullify any effect of cell density on nitrite production. ***p<0.001

PPI network for upregulated genes in *Panchvalkal-*exposed *P. aeruginosa* generated through STRING is presented in Figure 5, which shows 121 nodes connected through 70 edges with an average node degree of 1.16. Though empirically the centrality of the upregulated genes appeared to be lesser than those downregulated in *Panchvalkal*-exposed *P. aeruginosa*; since the number of edges (70) in this PPI network is 1.89-fold higher than expected (37) with a PPI enrichment p-value 1.27e-06, this network can be said to possess significantly more interactions among the member proteins than what can be expected for a random set of proteins of this much sample size and degree distribution. Such an enrichment can be taken as an indication of the member proteins being at least partially biologically connected. When we arranged the 121 upregulated genes in decreasing order of node degree, 62 nodes were found to have a non-zero score, and we selected top 26 genes with a node degree ≥3 (Table S5) for further ranking by different cytoHubba methods. Then we looked for genes which appeared among top ranked candidates by ≥6 cytoHubba methods, and 14 such genes (Table S6) were identified for further cluster analysis. Interaction map of these 14 important genes (Figure 6) showed them to be networked with the average node degree score of 2.29. Number of edges possessed by this network was 16 as against expected 1 for any such random set of proteins. These 14 genes were found to be distributed among five different local network clusters. Strength score for each of these clusters was >1.5. While three of the proteins (*atsB, msuE*, and *ssuB1*) were common members of 3 different clusters, one gene (*tauA*) appeared in two clusters. All these four highly networked upregulated genes (Table 2) are involved in sulfur metabolism in *P. aeruginosa* [Tralau et al., 2007]. Hence it may be speculated that *Panchvalkal* has induced sulfur starvation in *P. aeruginosa*, to overcome which the pathogen is forced to upregulate genes involved in sulfur transport and metabolism.

**Table 2.** Hubs identified as potential targets from among the up-regulated genes in *Panchvalkal*-exposed *P. aeruginosa*.

| No. | Gene ID/name | Codes for | Remarks |
| --- | --- | --- | --- |
| 1 | PA2357/<br>*msuE* (*slfA*) | FMN reductase | Involved in Riboflavin metabolism and sulfur metabolism pathways |
| 2 | PA3442/<br>*ssub1* | Aliphatic sulfonates import ATP-binding protein SsuB 1 | Aliphatic sulfonates import ATP-binding protein SsuB 1; Part of the ABC transporter complex SsuABC involved in aliphatic sulfonates import. Responsible for energy coupling to the transport system. |
| 3 | PA0185/*atsB* | Serine-modifying enzyme (Marquardt et al., 2003); Probable permease of ABC transporter | *atsB* is a member of a *cys* regulon in *P. aeruginosa*, which constitutes a general sulfate ester transport system (Hummerjohann et al., 2000) |
| 4 | PA3938/*tauA* | TauA (Sulfonate transport system ATP-binding protein); | This probable periplasmic taurine-binding protein precursor is part of *tau* operon involved in sulfur metabolism. |

**Figure 5.**
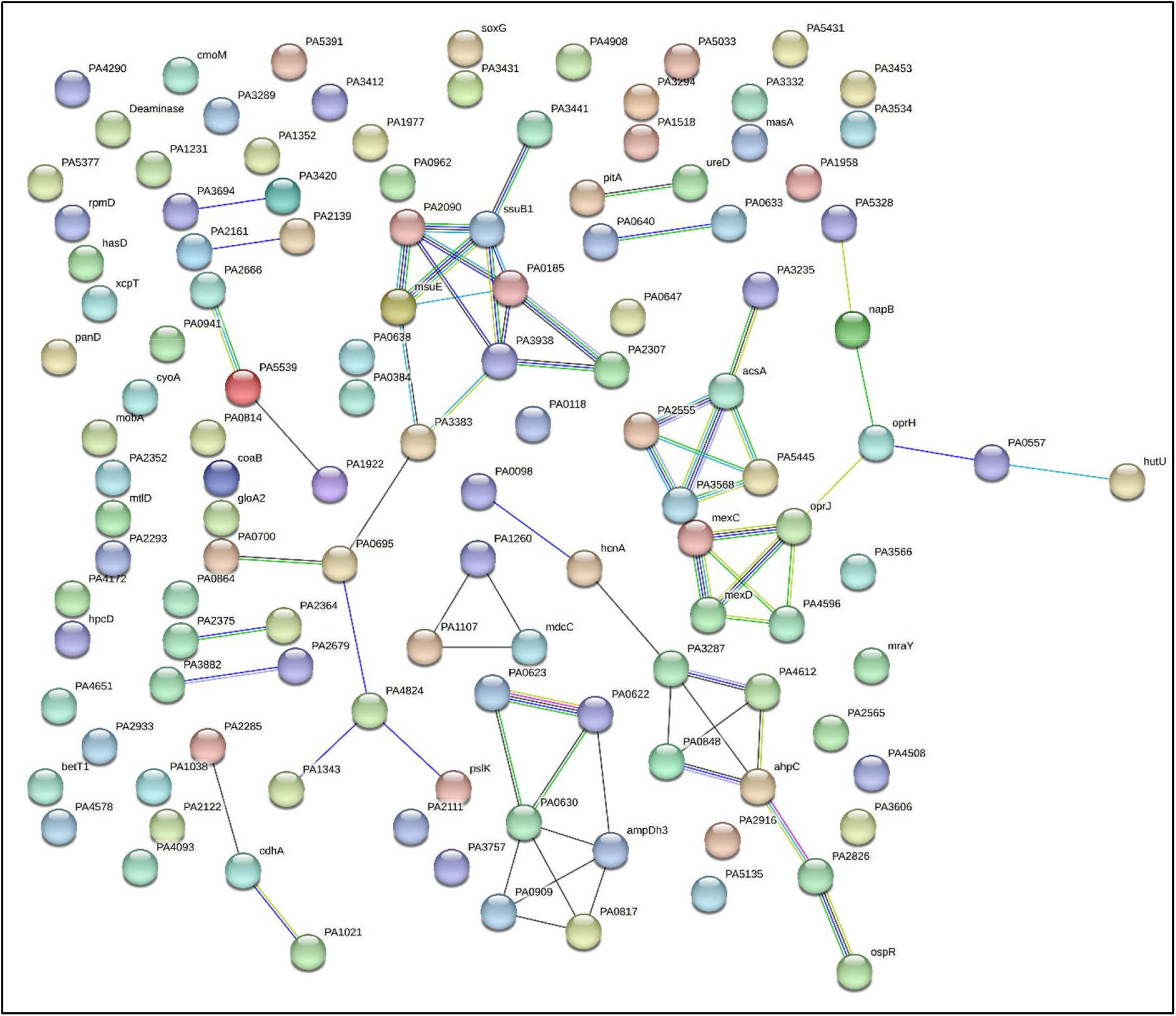
Protein-Protein Interaction (PPI) network of up-regulated genes in *Panchvalkal*-exposed *P. aeruginosa*.

**Figure 6.**
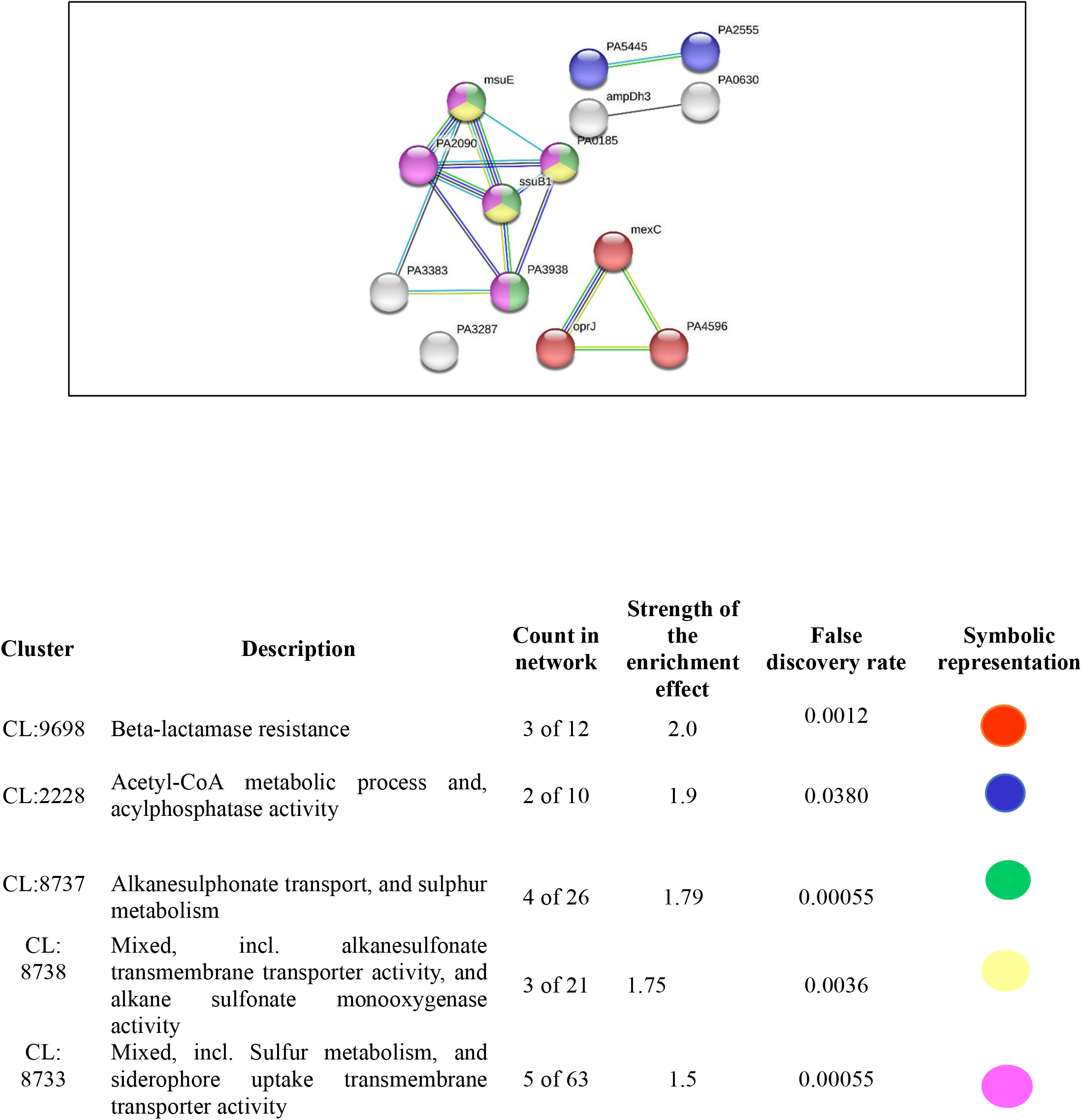
PPI network of top-ranked genes revealed through cytoHubba among up-regulated DEG in *Panchvalkal*-exposed *P. aeruginosa*.

### Network analysis of DEG in Herboheal*-*exposed *S. aureus*

Herboheal is a folk-inspired wound-healing formulation, and we had earlier demonstrated its anti-virulence potential against multiple bacterial pathogens including *Staphylococcus aureus*. Pre-treatment of *S. aureus* with Herboheal (0.1%v/v) could attenuate its virulence towards the surrogate host *C. elegans* by 55%. This concentration had a moderate growth-inhibitory effect (32%) on *S. aureus*, while heavily inhibiting staphyloxanthin production (79%). Whole transcriptome study revealed that approximately 17% of the *S. aureus* genome was expressed differently under the influence of Herboheal. The total number of DEG satisfying the dual criteria of log fold-change ≥2 and FDR≤0.01 was 113, of which 57 were up-regulated and 56 were downregulated (Table S7). Since the number of genes amenable to mapping by STRING turned out to be only 28 of these 113, we went for a combined PPI network (Figure 7) of these all DEG instead of preparing separate PPI map of upregulated or downregulated genes. The said PPI network had 28 nodes connected through 36 edges with an average node degree of 2.57. Since the number of edges (36) in this PPI network is 3-fold higher than expected (12) with a PPI enrichment p–value 1.02e-08, this network can be said to possess significantly more interactions among the member proteins than what can be expected for a random set of proteins having identical sample size and degree distribution. Such an enrichment is suggestive of the member proteins being at least partially biologically connected.

**Figure 7.**
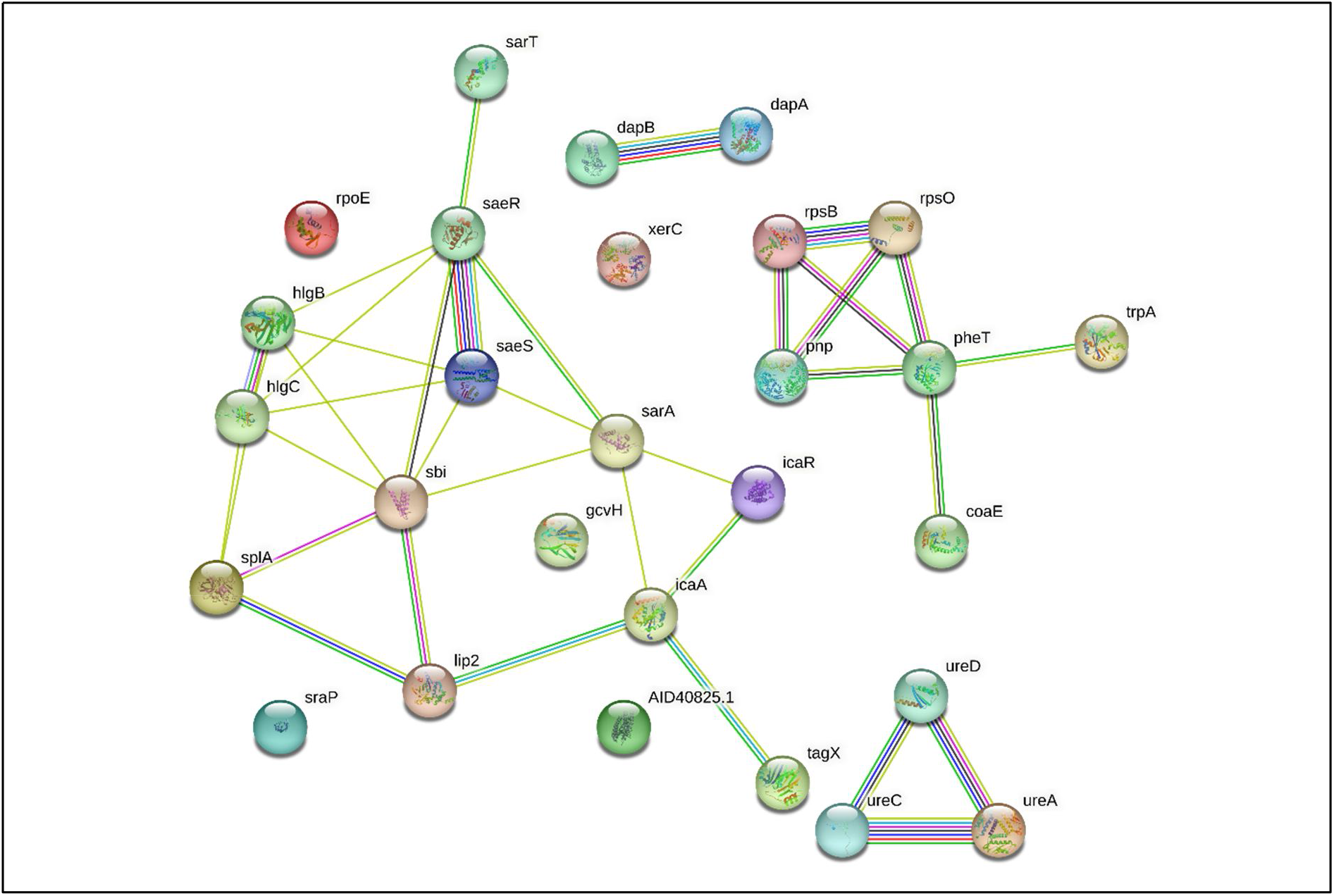
PPI network of up-down regulated genes in Herboheal*-*exposed *S. aureus*.

When we arranged all the 28 nodes in decreasing order of node degree, 23 nodes were found to have a non-zero score, and we selected top 13 genes with a node degree ≥3 (Table S8) for further ranking by different cytoHubba methods. Then we looked for genes which appeared among top ranked candidates by ≥6 cytoHubba methods. Of such 12 genes, 8 (Table S9) which were ranked among top-10 by ≥11 cytoHubba methods were taken for further cluster analysis. Interaction map of these 8 important genes (Figure 8) showed them to be networked with the average node degree score of 4. Number of edges possessed by this network was 16 as against expected 1 for any such random set of proteins. These 8 genes were found to be distributed among three different local network clusters. Strength score for each of these clusters was >1.46. While three of the proteins (*sarA, sbi* and *splA*) were common members of 2 different clusters. Four proteins were part of any one cluster, while *pnp* was not shown to be connected to remaining 7 genes. Since in case of *S. aureus*, we analysed up- and down-regulated genes together, instead of considering only the multi-cluster proteins as hubs, we took all of those which appeared to be part of PPI network shown in Figure-7. Functions of these 7 potential hubs are listed in Table-3.

**Figure 8.**
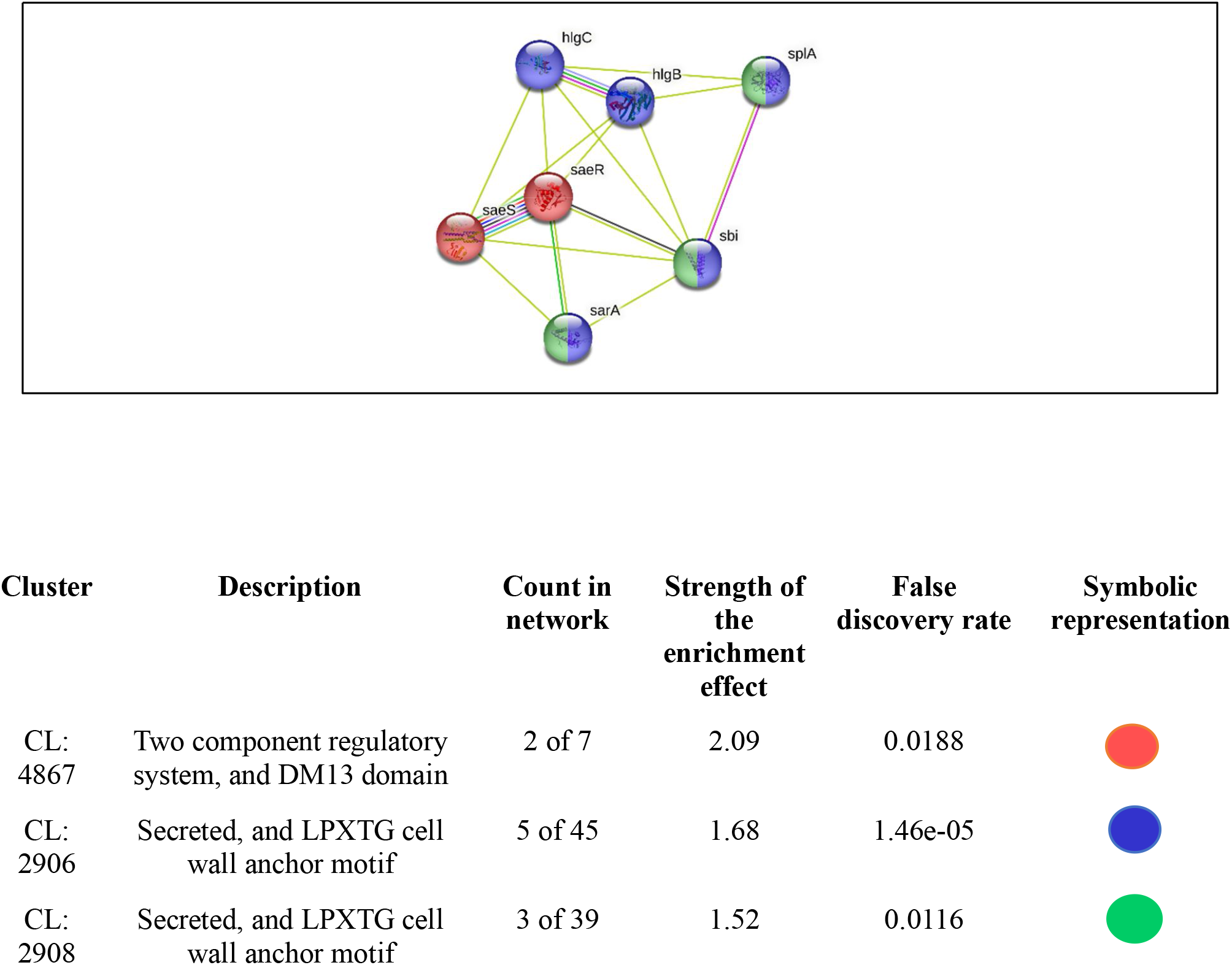
PPI network of top-ranked genes revealed through cytoHubba among down-regulated DEG Herboheal*-*exposed *S. aureus*.

## Discussion

*Panchvalkal*-exposed *P. aeruginosa* appears to suffer from sulfur-starvation and nitrosative stress. Compromised nitric oxide (NO) detoxification can render bacteria more susceptible to the NO produced by the host immune system (Arai, 2011). Mutant *P. aeruginosa* deficient in NO reductase was shown to register a reduced survival rate in NO-producing macrophages (Kakishima et al., 2007). NO has a strategic role in the metabolism of microorganisms in natural environments and also during host–pathogen interactions. NO as a signalling molecule is able to influence group behaviour in microorganisms. Downregulation of the denitrification pathway can disturb the homeostasis of the bacterial biofilms. NO levels can also affect motility, attachment and group behaviour in bacteria by affecting various signalling pathways involved in the metabolism of 3’,5’-cyclic di-guanylic acid (c-di-GMP). Suppressing bacterial detoxification of NO can be an effective anti-pathogenic strategy, as NO is known to modulate several aspects of bacterial physiology, including protection from oxidative stress and antimicrobials, homeostasis of the bacterial biofilm, etc (Gusarov and Nudler, 2005; Gusarov et al., 2009; Rinaldo et al., 2018). From this *in silico* exercise, nitric oxide reductase (NOR) has emerged as the most important target of *Panchvalkal* in *P. aeruginosa*. NOR is one of the important detoxifying enzymes of this pathogen, which is crucial to its ability to withstand nitrosative stress, and has also been reported to be important for virulence expression of this pathogen, and thus can be a plausible potential target for novel anti-virulence agents. (Arai, 2011). NOR inhibitors can be expected to compromise the pathogen’s ability to detoxify nitric oxide (NO), not allowing its virulence traits (e.g., biofilm formation, as NO has been indicated to act as a biofilm-dispersal signal) to be expressed fully. NOR inhibitors can be expected to be effective not only against *P. aeruginosa* but against other multiple pathogens too, as NO is reported to be perceived as a dispersal signal by various gram-negative and gram-positive bacteria (Barraud et al., 2015). This is to say, NOR inhibitors may be expected to have a broad spectrum activity against multiple pathogens. Major function of NOR is to detoxify NO generated by nitrite reductase (NIR). NO is a toxic byproduct of anaerobic respiration in *P. aeruginosa*. NO-derived nitrosative species can damage DNA, and compromise protein function. Intracellular accumulation of NO is likely to be lethal for the pathogen. It can be logically anticipated that *P. aeruginosa*’s ability to detoxify NO will be compromised under the influence of potent NOR inhibitors like *Panchvalkal*. Since NO seems to have a broad-spectrum anti-biofilm effect, NOR activity is essential for effective biofilm formation by the pathogens. NOR activity and NO concentration can modulate cellular levels of c-di-GMP, which is a secondary messenger molecule recognized as a key bacterial regulator of multiple processes such as virulence, differentiation, and biofilm formation. (Plate and Marletta, 2012). In the mammalian pathogens, the host’s macrophages are a likely source of NO. NOR expressed by the pathogen provides protection against the host defense mechanism (Kakishima et al., 2007). Since NOR activity is known to be important in multiple pathogenic bacteria (e.g., *P. aeruginosa, Staphylococcus aureus, Serratia marcescens*) for biofilm formation, virulence expression, combating nitrosative stress, and evading hose defense; NOR seems to be an important target for novel broad spectrum anti-pathogenic agents. A potential NOR inhibitor besides troubling the pathogen directly, may also boost its clearance by the host macrophages. (Schairer et al., 2012)

Based on the analysis of differently expressed upregulated genes, sulfur-starved culture of *P. aeruginosa* can be expected to experience compromised virulence. Upregulation of organic sulfur transport and metabolism genes has been reported in *P. aeruginosa* facing sodium hypochlorite-induced oxidative stress (Small et al., 2007a). Two of the upregulated hubs mentioned in Table-2 are part of tau or ssu gene clusters, which are reported in gram-negative bacteria like Escherichia coli too for being necessary for the utilization of taurine and alkanesulfonates as sulfur sources. Since these genes are exclusively expressed under conditions of sulfate or cysteine starvation (Eichhorn et al., 2000), one of the multiple effects exerted by Panchvalkal on *P. aeruginosa* can be said to be sulfur starvation. Upregulation of n-alkanesulfonates or taurine (sources of carbon and organic sulfur) utilization genes in *P. aeruginosa* suggests that the sulfur in these compounds was used to counter *Panchvalkal*-induced sulfur starvation, and that the neutrophilic amines and alpha-amino acids formed by catabolization of n-alkanesulfonates may guard the cell against oxidative stress (Small et al., 2007b). Thus, depriving *P. aeruginosa* of sulfur can be viewed as a potential anti-virulence strategy.

Among the potential targets identified in *S. aureus* in this study, first we discuss two such down-regulated genes which are common members of two different clusters. Of them, *splA* is a serine protease, exclusively specific of *S. aureus*, and thought to have a role in the second invasive stage of the infection (Stec-Niemczyk et al., 2009). Another potential hub *sbi* is an IgG-binding protein, which has a role in the inhibition of the innate as well as adaptive immune responses. Its secreted form acts as a potent complement inhibitor of the alternative pathway-mediated lysis. *sbi* helps mediate bacterial evasion of complement via a mechanism called futile fluid-phase consumption (Burman et al., 2008). Among the remaining potential hubs listed in Table-3, *SaeR/S* two component system is recognized as a major contributor to *S. aureus* pathogenesis and neutrophil evasion. The *SaeR/S* also plays role in regulating such virulence factors which decrease neutrophil hydrogen peroxide and hypochlorous acid production following *S. aureus* phagocytosis (Guerra et al., 2016). *S. aureus’s* escapes from the antimicrobial proteins NETs (neutrophil extracellular traps) is dependent on its secreting nuclease (*nuc*), and the latter in turn is regulated by *SaeR/S*. The *SaeR/S* system also modulates neutrophil fate by inhibiting IL-8 production and NF-κB activation. *saeR/S* deletion mutant of *S. aureus* was shown to be inferior than its wild-type counterpart in causing programmed neutrophil death (Guerra et al., 2017). The *SaeR/S* system regulates expression of many important virulence factors in *S. aureus*, and some of them do appear in our list of important targets such as *sbi, hlgB* and *hlgC*. Thus, inhibiting *SaeR/S* from sensing its environment can be expected to prevent expression of a multitude of *S. aureus* virulence factors in response to host signals. *hlgB* and *hlgC* are haemolytic proteins, and such proteins are used by many pathogens to fulfil their iron requirement as the concentration of free iron in human serum is much lesser than that required by the bacteria (Wilson, 2005). Down-regulation of bacterial haemolytic machinery may push them towards iron-starvation and thus compromising their fitness for in-host survival. Among all the potential hubs identified in Herboheal-exposed *S. aureus*, only one (*sarA)* was upregulated, and its upregulation seems to be a response from *S. aureus* to compensate the Herboheal-induced down-regulation of many important virulence traits. For example, *sarA* regulates expression of *ica* operon, which is required for biofilm formation in *S. aureus*. It can be said that *S. aureus*’s ability to adhere to surfaces and biofilm formation was compromised in presence of Herboheal as suggested by downregulation of adhesion/biofilm-relevant genes (*SaeR/S* and *sarA*), and as an adaptation to such challenge the pathogen is trying to upregulate *SarA*. This corroborates well with our previous report describing 56% reduced biofilm formation by *S. aureus* in presence of Herboheal (Patel et al., 2019).

This study has identified certain potential hubs in *P. aeruginosa* (Table 1-2) and *S. aureus* (Table 3) which should further be investigated for their candidature as potential anti-pathogenic targets. The most suitable targets in bacterial pathogens would be the ones which are absent from their host, as this will allow the criteria of selective toxicity to be satisfied for a newly discovered drug. We did a gene cooccurrence pattern analysis of gene families across genomes (through STRING) with respect to the major hubs identified in each of the pathogen (Table 4). Of the 19 hubs identified in either of the pathogen, none was shown to be present in *Homo sapiens*, and hence drugs causing dysregulation of one or more of these genes in pathogens are less likely to be toxic to humans.

**Table 3.** Hubs identified as potential targets from among the up and down-regulated genes in Herboheal*-*exposed *S. aureus*.

| No. | Gene ID | Gene Name | Codes for | Function |
| --- | --- | --- | --- | --- |
| 1 | 1280.SAXN108_0683 | *sarA* | Transcriptional regulator SarA | Probably activates the development of biofilm by both enhancing the *ica* operon transcription and suppressing the transcription of either a protein involved in the turnover of PIA/PNAG or a repressor of its synthesis, whose expression would be sigma-B-dependent. |
| 2 | 1280.SAXN108_2673 | *sbi* | Immunoglobulin-binding protein Sbi | Plays a role in the inhibition of both the innate and adaptive immune responses |
| 3 | 1280.SAXN108_1846 | *splA* | Serine protease SplA | Poorly characterized secreted protein probably involved in virulence |
| 4 | 1280.SAXN108_0774 | *saeR* | Response regulator transcription factor SaeR | The *saeR/S* system plays role in regulating such virulence factors which decrease neutrophil hydrogen peroxide and hypochlorous acid production following *S. aureus* phagocytosis |
| 5 | 1280.SAXN108_0773 | *saeS* | Histidine kinase |  |
| 6 | 1280.SAXN108_2677 | *hlgB* | Gamma-hemolysin component B precursor | Toxins that seems to act by forming pores in the membrane of the cell; Has a hemolytic and a leucotoxic activity. |
| 7 | 1280.SAXN108_2676 | *hlgC* | Gamma-hemolysin component C precursor |  |

**Table 4.**
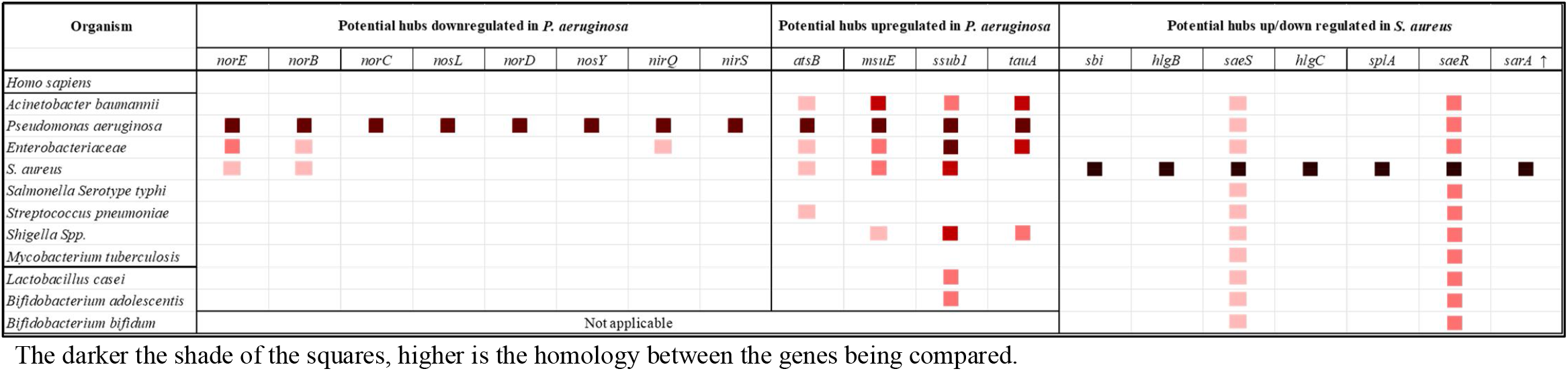
Cooccurrence analysis of genes coding for potential targets in *P. aeruginosa* and *S. aureus*.

If any target gene is present among multiple pathogens, then it can be considered suitable for a broad-spectrum antibacterial. We analysed the cooccurrence of identified hubs among some of the important pathogens listed by CDC and WHO. From among those listed in Table-4, *atsB, msuE, ssub1, norE*, and *norB* seemed to be present in multiple gram-negative as well as gram-positive pathogens, and thus suitable to be targeted by a broad-spectrum anti-pathogenic discovery programme. On the other hand, *tauA* and *nirQ* seemed to be present only among gram-negative pathogens. They can prove important targets in light of the fact that discovery of novel antimicrobials against gram-negative bacteria is relatively more challenging (Muñoz and Hergenrother, 2021).

One of the issues with conventional antibiotics is that they cannot differentiate between the ‘good’ (symbionts in human microbiome) and ‘bad’ (pathogens) bacteria, and hence their consumption may lead to gut dysbiosis. An ideal antimicrobial agent should target pathogens exclusively without causing gut dysbiosis. In this respect, a target in pathogenic bacteria absent from symbionts of human microbiome will be most suitable candidate for antibiotic discovery programmes. To gain some insight on this front regarding the targets identified by us, we run a gene cooccurrence analysis with some representative ‘good’ bacteria reported to be part of healthy human microbiome. *Bifidobacterium* species showed presence of no other target except *SaeR/S. SaeR/S* being widely distributed among bacteria can be considered a valid target, however an antibacterial agent targeting it may lead to gut dysbiosis too. All downregulated targets in P. aeruginosa were absent from the selected symbionts, which further adds value to their potential candidature as anti-virulence targets. However *atsB* and *ssub1* appeared to be present in *Lactobacillus casei*.

## Conclusion

This study has identified certain potential targets in two important pathogens. Further work is warranted on validation of the identified targets. Deletion mutants of the identified hub genes should be assessed for their expected attenuated virulence in appropriate host models. Next-generation of pathoblockers targeting any one of these genes may not always be effective as standalone therapeutic, and simultaneous targeting of more than one of these genes may be required for an effective therapy. They can also prove to be useful adjuvants to conventional antibiotics allowing use of bactericidal antibiotics at lower concentrations.

Besides indicating generation of nitrosative stress, inducing sulfur starvation, and disturbing regulation of bacterial virulence as potentially effective anti-pathogenic strategies, this study also demonstrates the relevance of the polyherbalism concept of the Traditional Medicine systems, and utility of the network analysis approach in elucidating the multiple modes of anti-pathogenic action exerted by the multicomponent natural extracts.

## Acknowledgement

Authors thank Nirma Education and Research Foundation (NERF), Ahmedabad for infrastructural support; Dr. Palep’s Medical Education and Research Foundation for providing *Panchvalkal* extract; Pooja Patel and Chinmayi Joshi for help with mining raw data.

## Abbreviations

AMR: Antimicrobial Resistance
DEG: Differentially Expressed Genes
NO: Nitric Oxide
NOR: Nitric Oxide Reductase
PPI: Protein-Protein Interaction

