## Supplementary File for "Network analysis for identifying potential anti-virulence targets through whole transcriptome analysis of *Pseudomonas aeruginosa* and *Staphylococcus aureus* exposed to certain anti-pathogenic polyherbal formulations"

**Table S1. List of down regulated genes in *Panchvalkal* exposed *P. aeruginosa* satisfying the dual criteria of log fold-change ≥2 and FDR≤0.01**

| No. | Feature ID/ Gene | Codes for | log fold change | FDR |
|---|---|---|---|---|
| 1 | PA0521 | Nitric oxide reductase NorE protein | 8.76 | 2.34E-10 |
| 2 | PA4962 | Inner membrane protein | 8.40 | 0.0001 |
| 3 | PA2182 | Hypothetical protein | 7.75 | 0.001 |
| 4 | PA2607 | tRNA 2-thiouridine synthesizing protein B | 6.75 | 0.003 |
| 5 | PA2980 | Hypothetical protein | 5.88 | 7.77E-05 |
| 6 | *norB* | Nitric oxide reductase subunit B | 5.44 | 0 |
| 7 | *norC* | Nitric oxide reductase subunit B | 5.04 | 0 |
| 8 | PA1827 | 3-oxoacyl-[acyl-carrier protein] reductase | 4.52 | 3.77E-06 |
| 9 | PA1013.1 | tRNA-Ser | 4.50 | 0.002 |
| 10 | *atuE* | Isohexenylglutaconyl-CoA hydratase | 4.50 | 0.009 |
| 11 | PA1492 | Hypothetical protein | 4.20 | 0.001 |
| 12 | PA2085 | Ring-hydroxylating dioxygenase small subunit | 4.16 | 0.01 |
| 13 | *nosL* | Copper chaperone NosL | 4.15 | 1.82E-05 |
| 14 | PA0525 | Nitric oxide reductase NorD protein | 4.14 | 0 |
| 15 | PA5071 | 16S ribosomal RNA methyltransferase RsmE | 3.57 | 0.001 |
| 16 | PA2146 | Hypothetical protein | 3.53 | 0.001 |
| 17 | *nosY* | Cu-processing system permease protein | 3.46 | 0.002 |
| 18 | PA3377 | Alpha-D-ribose 1-methylphosphonate 5-phosphate C-P lyase | 3.45 | 0.0002 |
| 19 | PA1211 | Hypothetical protein | 3.40 | 0.01 |
| 20 | PA0818 | Hypothetical protein | 3.30 | 0.004 |
| 21 | PA3033 | Hypothetical protein | 3.25 | 0.0005 |
| 22 | *kynB* | Arylformamidase (kynurenine formamidase) | 3.15 | 0.0001 |
| 23 | PA5196 | Hypothetical protein | 3.14 | 0.005 |
| 24 | PA5181.1 | P34 | 3.11 | 0.001 |
| 25 | *nuoI* | NADH-quinone oxidoreductase subunit I | 3.08 | 2.73E-08 |
| 26 | PA4702 | Hypothetical protein | 3.04 | 1.11E-09 |
| 27 | *algE* | Alginate production protein | 3.00 | 0.003 |
| 28 | PA0526 | Hypothetical protein | 2.95 | 0 |
| 29 | PA2180 | Hypothetical protein | 2.94 | 0.004 |
| 30 | *nirQ* | Nitric oxide reductase NorQ protein | 2.91 | 0 |
| 31 | PA3493 | Electron transport complex protein RnfG \| | 2.80 | 3.64E-05 |
| 32 | *nirS* | Heme d1 biosynthesis protein | 2.78 | 0 |
| 33 | *infA* | Translation initiation factor IF-1 | 2.73 | 4.90E-06 |
| 34 | PA1879 | Hypothetical protein | 2.71 | 1.94E-05 |
| 35 | PA0270 | Hypothetical protein | 2.70 | 0.002 |

| No. | Feature ID/ Gene | Codes for | log fold change | FDR |
|---|---|---|---|---|
| 36 | PA4466 | Phosphoryl carrier protein | 2.68 | 9.20E-06 |
| 37 | PA0806 | Hypothetical protein | 2.68 | 0.006 |
| 38 | *rluA* | Ribosomal large subunit pseudouridine synthase A | 2.66 | 0.001 |
| 39 | PA2506 | Hypothetical protein | 2.66 | 0.01 |
| 40 | *lldD* | L-lactate dehydrogenase | 2.65 | 4.74E-05 |
| 41 | PA2570.1 | tRNA-Leu | 2.63 | 0.008 |
| 42 | PA2433 | Hypothetical protein | 2.62 | 1.35E-09 |
| 43 | *pcaC* | 4-carboxymuconolactone decarboxylase | 2.62 | 0.001 |
| 44 | *pilE* | Type IV pilus assembly protein | 2.62 | 0.001 |
| 45 | PA2754a | Hypothetical protein | 2.59 | 0.005 |
| 46 | PA0682 | HxcX atypical pseudopilin | 2.55 | 0.001 |
| 47 | *pscL* | Type III secretion protein L | 2.55 | 0.01 |
| 48 | PA2602 | Hypothetical protein | 2.54 | 0.006 |
| 49 | *pcaH* | Protocatechuate 3,4-dioxygenase | 2.54 | 0.006 |
| 50 | PA3580 | Cys-tRNA(Pro)/Cys-tRNA(Cys) deacylase | 2.48 | 5.39E-09 |
| 51 | PA5535 | Hypothetical protein | 2.44 | 3.13E-08 |
| 52 | PA0179 | Two-component system, chemotaxis family, response regulator CheY | 2.43 | 5.44E-06 |
| 53 | PA1312 | Transcriptional regulator | 2.42 | 0.0005 |
| 54 | PA0431 | Hypothetical protein | 2.42 | 0.005 |
| 55 | PA2039 | Hypothetical protein | 2.42 | 0.01 |
| 56 | PA5115 | Hypothetical protein | 2.40 | 0.005 |
| 57 | *gntR* | Transcriptional regulator GntR | 2.37 | 4.44E-16 |
| 58 | PA2162 | (1->4)-alpha-D-glucan 1-alpha-D-glucosylmutase | 2.37 | 1.51E-05 |
| 59 | PA3274 | Hypothetical protein | 2.36 | 9.66E-05 |
| 60 | PA3275 | Small multidrug resistance family-3 protein | 2.36 | 0.001 |
| 61 | PA2150 | DNA end-binding protein Ku | 2.35 | 0.0006 |
| 62 | *narH* | Respiratory nitrate reductase beta chain | 2.34 | 2.12E-05 |
| 63 | PA2136 | Hypothetical protein | 2.33 | 0.0003 |
| 64 | *nadE* | NH3-dependent NAD synthetase | 2.32 | 0.004 |
| 65 | PA4171 | Protease I | 2.31 | 0.003 |
| 66 | PA0924 | Hypothetical protein | 2.29 | 1.81E-09 |
| 67 | PA3880 | Hypothetical protein | 2.28 | 0.0007 |
| 68 | PA0544 | Hypothetical protein | 2.27 | 2.71E-08 |
| 69 | PA3450 | Antioxidant protein | 2.25 | 0.006 |
| 70 | PA0522 | Hypothetical protein | 2.25 | 0.01 |
| 71 | PA0952 | Hypothetical protein | 2.20 | 7.23E-05 |
| 72 | PA5155 | Polar amino acid transport system permease protein | 2.20 | 0.01 |
| 73 | PA0830 | Hypothetical protein | 2.19 | 2.18E-14 |
| 74 | PA1093 | Flagellar protein FlaG | 2.19 | 0.008551 |
| 75 | PA4064 | Putative ABC transport system ATP-binding protein | 2.18 | 0.007 |
| 76 | *cueR* | Copper efflux regulator | 2.15 | 0.0002 |
| 77 | PA0942 | Transcriptional regulator | 2.14 | 0 |
| 78 | PA0515 | Heme d1 biosynthesis protein NirD | 2.14 | 8.23E-05 |
| 79 | PA1020 | Acyl-CoA dehydrogenase | 2.14 | 0.007 |
| 80 | PA4921 | Hypothetical protein | 2.12 | 0.0001 |
| 81 | PA1763 | Hypothetical protein | 2.12 | 0.001 |
| 82 | *glpD* | Glycerol-3-phosphate dehydrogenase | 2.10 | 1.14E-12 |
| 83 | PA0121 | Hypothetical protein | 2.10 | 2.00E-05 |
| 84 | PA3172 | Phosphoglycolate phosphatase | 2.09 | 6.52E-05 |

| No. | Feature ID/ Gene | Codes for | log fold change | FDR |
|---|---|---|---|---|
| 85 | PA3859 | Phospholipase/carboxylesterase | 2.09 | 0.0005 |
| 86 | *moeA1* | Molybdopterin molybdotransferase | 2.08 | 1.48E-06 |
| 87 | *braZ* | Branched-chain amino acid:cation transporter | 2.08 | 0.0002 |
| 88 | PA0218 | Transcriptional regulator | 2.08 | 0.0003 |
| 89 | PA3016 | Hypothetical protein | 2.08 | 0.0009 |
| 90 | PA4357 | Ferrous iron transport protein C | 2.06 | 1.11E-12 |
| 91 | PA3224 | Hypothetical protein | 2.06 | 6.11E-07 |
| 92 | PA1221 | Hypothetical protein | 2.06 | 0.002 |
| 93 | PA3459 | Asparagine synthase | 2.05 | 0 |
| 94 | PA0665 | Iron-sulfur cluster insertion protein | 2.05 | 9.64E-12 |
| 95 | PA0443 | Nucleobase:cation symporter-1, NCS1 family | 2.05 | 0.01 |
| 96 | PA0828 | Transcriptional regulator | 2.04 | 0.0009 |
| 97 | PA1015 | Transcriptional regulator | 2.04 | 0.004 |
| 98 | PA3913 | Putative protease | 2.03 | 1.03E-05 |
| 99 | PA1057 | Multicomponent K+:H+ antiporter subunit E | 2.03 | 0.01 |
| 100 | *ohrR* | Transcriptional regulator | 2.03 | 0.01 |
| 101 | PA0177 | Purine-binding chemotaxis protein CheW | 2.03 | 0.01 |
| 102 | PA3847 | Hypothetical protein | 2.03 | 0.01 |
| 103 | PA1470 | 3-oxoacyl-[acyl-carrier protein] reductase | 2.02 | 0.003 |
| 104 | PA0911 | Hypothetical protein | 2.02 | 0.01 |
| 105 | PA3070 | MoxR-like ATPase | 2.01 | 5.10E-07 |

Genes are arranged in decreasing order of Fold Change.

**Table S2. Node degree score of the genes mentioned in Table S1**

| No. | Gene ID/ Symbol | Identifier | Node degree |
|---|---|---|---|
| 1 | *nirS* | 208964.PA0519 | 11 |
| 2 | *norB* | 208964.PA0524 | 11 |
| 3 | PA0525 | 208964.PA0525 | 10 |
| 4 | *nirQ* | 208964.PA0520 | 10 |
| 5 | *norC* | 208964.PA0523 | 10 |
| 6 | *nosL* | 208964.PA3396 | 10 |
| 7 | PA0521 | 208964.PA0521 | 9 |
| 8 | *nosY* | 208964.PA3395 | 9 |
| 9 | PA3913 | 208964.PA3913 | 8 |
| 10 | PA0515 | 208964.PA0515 | 6 |
| 11 | PA0522 | 208964.PA0522 | 6 |
| 12 | PA2146 | 208964.PA2146 | 6 |
| 13 | *narH* | 208964.PA3874 | 6 |
| 14 | *anvM* | 208964.PA3880 | 5 |
| 15 | PA0177 | 208964.PA0177 | 3 |
| 16 | PA1763 | 208964.PA1763 | 3 |
| 17 | PA2180 | 208964.PA2180 | 3 |
| 18 | PA3274 | 208964.PA3274 | 3 |
| 19 | *ku* | 208964.PA2150 | 3 |
| 20 | *nadE* | 208964.PA4920 | 3 |
| 21 | PA0828 | 208964.PA0828 | 2 |
| 22 | *algE* | 208964.PA3544 | 2 |
| 23 | *moeA1* | 208964.PA3914 | 2 |
| 24 | *nuoI* | 208964.PA2644 | 2 |
| 25 | *pcaH* | 208964.PA0153 | 2 |
| 26 | PA0179 | 208964.PA0179 | 1 |
| 27 | PA0544 | 208964.PA0544 | 1 |
| 28 | PA0682 | 208964.PA0682 | 1 |
| 29 | PA0830 | 208964.PA0830 | 1 |
| 30 | PA0924 | 208964.PA0924 | 1 |
| 31 | PA0952 | 208964.PA0952 | 1 |
| 32 | PA1015 | 208964.PA1015 | 1 |
| 33 | PA1020 | 208964.PA1020 | 1 |
| 34 | PA1211 | 208964.PA1211 | 1 |
| 35 | PA1221 | 208964.PA1221 | 1 |
| 36 | PA1470 | 208964.PA1470 | 1 |
| 37 | PA1827 | 208964.PA1827 | 1 |
| 38 | PA2136 | 208964.PA2136 | 1 |
| 39 | PA2433 | 208964.PA2433 | 1 |
| 40 | PA3016 | 208964.PA3016 | 1 |
| 41 | PA3224 | 208964.PA3224 | 1 |
| 42 | PA3450 | 208964.PA3450 | 1 |
| 43 | PA3859 | 208964.PA3859 | 1 |
| 44 | PA5071 | 208964.PA5071 | 1 |
| 45 | *choE* | 208964.PA4921 | 1 |
| 46 | *erpA* | 208964.PA0665 | 1 |
| 47 | *glpD* | 208964.PA3584 | 1 |
| 48 | *infA* | 208964.PA2619 | 1 |
| 49 | *kynB* | 208964.PA2081 | 1 |
| 50 | *pcaC* | 208964.PA0232 | 1 |
| 51 | *pscL* | 208964.PA1725 | 1 |
| 52 | *rluA* | 208964.PA3246 | 1 |

Rest 48 genes with node degree score ‘zero’ are not listed.

**Table S3. Top ten cytoHubba ranked genes from among the top-13 in Table S2**

| No. | Gene ID | Gene Name | Number of methods ranking this protein among top 10 | Names of 12 ranking methods of CytoHubba and rank score provided by them | | | | | | | | | | | |
|---|---|---|---|---|---|---|---|---|---|---|---|---|---|---|---|
| | | | | Degree | MNC | DMNC | MCC | Bottleneck | EcCentricity | Closeness | Radiality | Betweenness | Stress | CC | EPC |
| 1 | PA0525 | *norD* | 12 | 10 | 10 | 0.71829 | 7200 | 2 | 0.325 | 11 | 1.35417 | 3.28571 | 18 | 0.8 | 5.793 |
| 2 | PA0520 | *nirQ* | 12 | 10 | 10 | 0.71829 | 7200 | 1 | 0.325 | 11 | 1.35417 | 3.28571 | 18 | 0.8 | 5.68 |
| 3 | PA3395 | *nosY* | 12 | 9 | 9 | 0.69213 | 5772 | 1 | 0.325 | 10.5 | 1.3 | 3.18571 | 14 | 0.80556 | 5.559 |
| 4 | PA3396 | *nosL* | 11 | 10 | 10 | 0.63848 | 5784 | 3 | 0.325 | 11 | 1.35417 | 7.01905 | 26 | - | 5.69 |
| 5 | PA0521 | *norE* | 11 | 9 | 9 | 0.73986 | 6480 | - | 0.325 | 10.5 | 1.3 | 1.66667 | 10 | 0.86111 | 5.492 |
| 6 | PA0519 | *nirS* | 10 | 11 | 11 | - | 6498 | 2 | 0.325 | 11.5 | 1.48033 | 13.4 | 38 | - | 5.84 |
| 7 | PA0524 | *norB* | 10 | 11 | 11 | 0.64479 | 7206 | - | 0.325 | 11.5 | 1.48033 | 9.11905 | 34 | - | 5.917 |
| 8 | PA0523 | *norC* | 9 | 10 | 10 | 0.71829 | 7200 | - | - | 11 | 1.35417 | 3.28571 | 18 | - | 5.572 |
| 9 | PA0522 | Hypothe-tical protein | 10 | 6 | 6 | 0.713237 | 720 | 1 | 0.325 | 9 | 1.1375 | | | 1 | 4.708 |
| 10 | PA3913 | UbiU | 10 | 7 | 7 | 0.621988 | 726 | - | - | 9.5 | 1.191667 | 1.9 | 8 | 0.809524 | 4.88 |

"-": This method did not rank the shown protein among top 10.

MNC: Maximum Neighborhood Component; DMNC: Density of Maximum Neighborhood Component; MCC: Maximal Clique Centrality; CC: Clustering Co-efficient; EPC: Edge Percolated Component

**Table S4. List of up regulated genes in *Panchvalkal* exposed *P. aeruginosa* satisfying the dual criteria of log fold-change ≥2 and FDR≤0.01**

| No. | Feature ID/Gene | Codes for | log fold change | FDR |
|---|---|---|---|---|
| 1 | *mexC* | Membrane fusion protein, multidrug efflux system | 16.82 | 0 |
| 2 | PA2139 | Pseudogene | 15 | 0.01 |
| 3 | PA3441 | Molybdopterin-binding protein | 10 | 0.006 |
| 4 | PA5328 | Mono-heme cytochrome C | 8 | 0.01 |
| 5 | PA2161 | Hypothetical protein | 7.22 | 4.16E-06 |
| 6 | *oprJ* | Outer membrane protein, multidrug efflux system | 6.91 | 0 |
| 7 | PA3383 | Phosphonate transport system substrate-binding protein | 6.33 | 0.0006 |
| 8 | PA0700 | Hypothetical protein | 5.8 | 0.003 |
| 9 | PA2565 | Hypothetical protein | 5.74 | 0 |
| 10 | PA0909 | Hypothetical protein | 5.4 | 0.005 |
| 11 | *napB* | Cytochrome c-type protein | 5.26 | 1.12E-13 |
| 12 | PA2090 | Hypothetical protein | 5.11 | 0.0004 |
| 13 | *hpcD* | 5-carboxymethyl-2-hydroxymuconate isomerase | 4.8 | 0.01 |
| 14 | PA2285 | Hypothetical protein | 4.8 | 1.55E-05 |
| 15 | PA3566 | Hypothetical protein | 4.75 | 0.001 |
| 16 | PA0695 | Hypothetical protein | 4.42 | 0.005 |
| 17 | PA1107 | Diguanylate cyclase | 4.21 | 0 |
| 18 | *mexD* | Multidrug efflux pump | 4.15 | 0 |
| 19 | PA2364 | Type VI secretion system protein | 4.03 | 1.29E-12 |
| 20 | *coaB* | Phage coat protein B | 4 | 0.01 |
| 21 | PA3442 | Sulfonate transport system ATP-binding protein | 4 | 0.004 |
| 22 | PA4866 | Phosphinothricin acetyltransferase | 4 | 0.002 |
| 23 | PA1231 | Hypothetical protein | 3.93 | 0.0002 |
| 24 | PA4172 | Exodeoxyribonuclease III | 3.93 | 5.00E-07 |
| 25 | PA0640 | Bacteriophage protein | 3.55 | 1.84E-07 |
| 26 | PA2499 | Deaminase | 3.53 | 0.002 |
| 27 | PA1352 | Hypothetical protein | 3.51 | 1.34E-05 |
| 28 | *ospR* | Transcriptional regulator | 3.43 | 0 |
| 29 | *cdhA* | Carnitine 3-dehydrogenase | 3.42 | 0.002 |
| 30 | PA4908 | Ornithine cyclodeaminase | 3.31 | 1.18E-06 |
| 31 | PA5391 | Hypothetical protein | 3.21 | 0.0008 |
| 32 | PA2307 | NitT/TauT family transport system permease protein | 3.18 | 8.69E-05 |
| 33 | PA5135 | Hypothetical protein | 2.97 | 9.42E-06 |
| 34 | PA2933 | large subunit ribosomal protein L6 (rplF; 50S ribosomal protein L6) | 2.96 | 0.0004 |
| 35 | PA3235 | Hypothetical protein | 2.96 | 1.62E-10 |
| 36 | PA4290 | Methyl-accepting chemotaxis protein | 2.96 | 0 |
| 37 | PA0384 | Hypothetical protein | 2.92 | 0.01 |
| 38 | PA3287 | Hypothetical protein | 2.92 | 5.20E-14 |

| No. | Feature ID/Gene | Codes for | log fold change | FDR |
|---|---|---|---|---|
| 39 | PA0848 | Peroxiredoxin (alkyl hydroperoxide reductase subunit C) | 2.9 | 2.08E-09 |
| 40 | PA1021 | enoyl-CoA hydratase | 2.88 | 0.0007 |
| 41 | PA2916 | Hypothetical protein | 2.86 | 0.009 |
| 42 | PA0638 | Bacteriophage protein | 2.84 | 8.11E-06 |
| 43 | PA0633 | Hypothetical protein | 2.84 | 1.29E-10 |
| 44 | *msuE* | FMN reductase | 2.75 | 0.01 |
| 45 | PA0623 | Bacteriophage protein | 2.74 | 9.84E-08 |
| 46 | *soxG* | Sarcosine oxidase | 2.73 | 0.002 |
| 47 | PA0814 | Hypothetical protein | 2.71 | 0.01 |
| 48 | PA2666 | 6-pyruvoyltetrahydropterin/6-carboxytetrahydropterin synthase | 2.7 | 0.009 |
| 49 | PA3431 | Hypothetical protein | 2.7 | 0.009 |
| 50 | PA5431 | GntR family transcriptional regulator | 2.67 | 1.41E-08 |
| 51 | PA0941 | Hypothetical protein | 2.66 | 0.01 |
| 52 | PA3757 | GntR family transcriptional regulator | 2.64 | 0.01 |
| 53 | PA2679 | Hypothetical protein | 2.61 | 0 |
| 54 | PA1343 | Bacteriophage protein | 2.59 | 1.03E-08 |
| 55 | *mraY* | Phospho-N-acetylmuramoyl-pentapeptide-transferase | 2.59 | 1.92E-10 |
| 56 | PA0622 | Bacteriophage protein | 2.59 | 3.12E-11 |
| 57 | PA0185 | Sulfonate transport system permease protein | 2.55 | 1.45E-06 |
| 58 | PA1260 | Polar amino acid transport system substrate-binding protein | 2.5 | 0.01 |
| 59 | PA4596 | Transcriptional regulator | 2.5 | 3.21E-09 |
| 60 | PA3606 | DTW domain-containing protein | 2.47 | 0.0001 |
| 61 | PA1977 | Hypothetical protein | 2.47 | 1.73E-05 |
| 62 | *hutU* | Urocanate hydratase | 2.46 | 0 |
| 63 | *mdcC* | Malonate decarboxylase delta subunit | 2.45 | 0.009 |
| 64 | *hasD* | ATP-binding cassette, subfamily C, bacterial exporter for protease/lipase | 2.45 | 5.11E-08 |
| 65 | *gloA2* | Lactoylglutathione lyase | 2.44 | 0.01 |
| 66 | *betT1* | Choline/glycine/proline betaine transport protein | 2.44 | 0.0009 |
| 67 | *pslK* | Polysaccharide biosynthesis protein PslK | 2.44 | 2.46E-05 |
| 68 | PA2122 | Hypothetical protein | 2.43 | 0.0002 |
| 69 | *ahpC* | Peroxiredoxin (alkyl hydroperoxide reductase subunit C) | 2.43 | 0 |
| 70 | PA3412 | Hypothetical protein | 2.42 | 0.01 |
| 71 | PA3938 | Taurine transport system substrate-binding protein | 2.41 | 0.002 |
| 72 | PA1038 | Hypothetical protein | 2.4 | 0.005 |
| 73 | PA1958 | Nicotinamide mononucleotide transporter | 2.39 | 0.0004 |
| 74 | PA5377 | Glycine betaine/proline transport system permease protein | 2.39 | 0.0002 |
| 75 | PA4093 | Hypothetical protein | 2.37 | 0.009 |
| 76 | PA1518 | 5-hydroxyisourate hydrolase | 2.36 | 0.01 |
| 77 | *ureD* | Urease accessory protein | 2.35 | 0.003 |
| 78 | PA0118 | Hypothetical protein | 2.33 | 0.01 |
| 79 | *mtlD* | Mannitol 2-dehydrogenase | 2.32 | 0.01 |

| No. | Feature ID/Gene | Codes for | log fold change | FDR |
|---|---|---|---|---|
| 81 | PA3453 | Hypothetical protein | 2.32 | 1.43E-07 |
| 82 | PA2352 | Glycerophosphoryl diester phosphodiesterase | 2.3 | 0.0008 |
| 83 | PA3534 | Oxidoreductase | 2.3 | 1.04E-06 |
| 84 | PA0962 | Starvation-inducible DNA-binding protein | 2.28 | 4.83E-09 |
| 85 | *hcnA* | Hydrogen cyanide synthase | 2.27 | 0.01 |
| 86 | *mobA* | Molybdenum cofactor guanylyltransferase | 2.27 | 0.01 |
| 87 | PA2111 | Hypothetical protein | 2.26 | 6.66E-16 |
| 88 | PA1922 | Outer membrane receptor for ferrienterochelin and colicins | 2.25 | 0.01 |
| 89 | PA4790 | S-adenosylmethionine-dependent methyltransferase | 2.25 | 0.006 |
| 90 | PA0647 | Hypothetical protein | 2.25 | 0.001 |
| 91 | PA4578 | Hypothetical protein | 2.25 | 6.67E-08 |
| 92 | PA4280.5 | 16S ribosomal RNA | 2.25 | 0 |
| 93 | *acsA* | Acetyl-CoA synthetase | 2.24 | 0 |
| 94 | PA4508 | Lrp/AsnC family transcriptional regulator, leucine-responsive regulatory protein | 2.22 | 0.002 |
| 95 | PA0098 | 3-oxoacyl-[acyl-carrier-protein] synthase I | 2.21 | 0.003 |
| 96 | PA4651 | Fimbrial chaperone protein | 2.19 | 0.001 |
| 97 | *xcpT* | Type II secretion system protein G | 2.17 | 0.002 |
| 98 | PA2555 | Acetyl-CoA synthetase | 2.17 | 8.88E-16 |
| 99 | PA0817 | Hypothetical protein | 2.16 | 0.01 |
| 100 | PA2375 | Hypothetical protein | 2.15 | 0.003 |
| 101 | *masA* | Enolase-phosphatase E1 | 2.15 | 2.30E-11 |
| 102 | PA2826 | Glutathione peroxidase | 2.14 | 1.89E-07 |
| 103 | PA5445 | Succinyl-CoA:acetate CoA-transferas | 2.13 | 4.02E-09 |
| 104 | PA0630 | Hypothetical protein | 2.11 | 0.009 |
| 105 | *panD* | Aspartate 1-decarboxylase | 2.1 | 0.006 |
| 106 | PA0557 | Hypothetical protein | 2.1 | 0.0009 |
| 107 | *cyoA* | Cytochrome o ubiquinol oxidase subunit II | 2.09 | 0.003 |
| 108 | PA0306 | Transcriptional regulator | 2.07 | 3.45E-05 |
| 109 | *oprH* | oprH; PhoP/Q and low Mg2+ inducible outer membrane protein H1 | 2.06 | 0 |
| 110 | PA2293 | Hypothetical protein | 2.05 | 0.007 |
| 111 | PA3694 | Hypothetical protein | 2.05 | 0.007 |
| 112 | PA3294 | Type VI secretion system secreted protein VgrG | 2.05 | 0.0002 |
| 113 | *rpmD* | Large subunit ribosomal protein L30 | 2.05 | 4.80E-09 |
| 114 | PA3289 | Hypothetical protein | 2.04 | 0.0009 |
| 115 | PA5539 | GTP cyclohydrolase I | 2.03 | 0.01 |
| 116 | PA0864 | Transcriptional regulator | 2.03 | 0.01 |
| 117 | PA3420 | Transcriptional regulator | 2.03 | 2.15E-06 |
| 118 | PA3568 | Propionyl-CoA synthetase | 2.03 | 2.01E-09 |
| 119 | *ampDh3* | N-acetylmuramoyl-L-alanine amidase | 2.02 | 0.008 |
| 120 | PA3332 | Hypothetical protein | 2 | 0.01 |
| 121 | PA3882 | Hypothetical protein | 2 | 0.006 |

| No. | Feature ID/Gene | Codes for | log fold change | FDR |
|---|---|---|---|---|
| 122 | PA4824 | Hypothetical protein | 2 | 0.002 |
| 123 | PA4612 | Hypothetical protein | 2 | 4.55E-07 |

Genes are arranged in decreasing order of Fold Change.

#### Table S5. Node degree score of the genes mentioned in Table S4

| No. | Gene ID / Symbol | Identifier | Node degree |
|---|---|---|---|
| 1 | PA0185 | 208964.PA0185 | 5 |
| 2 | PA0630 | 208964.PA0630 | 5 |
| 3 | PA3938 | 208964.PA3938 | 5 |
| 4 | *ssuB1* | 208964.PA3442 | 5 |
| 5 | PA2090 | 208964.PA2090 | 4 |
| 6 | PA3287 | 208964.PA3287 | 4 |
| 7 | *acsA* | 208964.PA0887 | 4 |
| 8 | *ahpC* | 208964.PA0139 | 4 |
| 9 | *ampDh3* | 208964.PA0807 | 4 |
| 10 | *msuE* | 208964.PA2357 | 4 |
| 11 | *oprJ* | 208964.PA4597 | 4 |
| 12 | PA0622 | 208964.PA0622 | 3 |
| 13 | PA0695 | 208964.PA0695 | 3 |
| 14 | PA0817 | 208964.PA0817 | 3 |
| 15 | PA0848 | 208964.PA0848 | 3 |
| 16 | PA0909 | 208964.PA0909 | 3 |
| 17 | PA2555 | 208964.PA2555 | 3 |
| 18 | PA3383 | 208964.PA3383 | 3 |
| 19 | PA3568 | 208964.PA3568 | 3 |
| 20 | PA4596 | 208964.PA4596 | 3 |
| 21 | PA4612 | 208964.PA4612 | 3 |
| 22 | PA4824 | 208964.PA4824 | 3 |
| 23 | PA5445 | 208964.PA5445 | 3 |
| 24 | *mexC* | 208964.PA4599 | 3 |
| 25 | *mexD* | 208964.PA4598 | 3 |
| 26 | *oprH* | 208964.PA1178 | 3 |
| 27 | PA0557 | 208964.PA0557 | 2 |
| 28 | PA0623 | 208964.PA0623 | 2 |
| 29 | PA1107 | 208964.PA1107 | 2 |
| 30 | PA1260 | 208964.PA1260 | 2 |

| No. | Gene ID / Symbol | Identifier | Node degree |
|---|---|---|---|
| 31 | PA2307 | 208964.PA2307 | 2 |
| 32 | PA2826 | 208964.PA2826 | 2 |
| 33 | PA5539 | 208964.PA5539 | 2 |
| 34 | *cdhA* | 208964.PA5386 | 2 |
| 35 | *hcnA* | 208964.PA2193 | 2 |
| 36 | *mdcC* | 208964.PA0210 | 2 |
| 37 | *napB* | 208964.PA1173 | 2 |
| 38 | PA0098 | 208964.PA0098 | 1 |
| 39 | PA0633 | 208964.PA0633 | 1 |
| 40 | PA0640 | 208964.PA0640 | 1 |
| 41 | PA0700 | 208964.PA0700 | 1 |
| 42 | PA1021 | 208964.PA1021 | 1 |
| 43 | PA1343 | 208964.PA1343 | 1 |
| 44 | PA1922 | 208964.PA1922 | 1 |
| 45 | PA2139 | 208964.PA2139 | 1 |
| 46 | PA2161 | 208964.PA2161 | 1 |
| 47 | PA2285 | 208964.PA2285 | 1 |
| 48 | PA2364 | 208964.PA2364 | 1 |
| 49 | PA2375 | 208964.PA2375 | 1 |
| 50 | PA2666 | 208964.PA2666 | 1 |
| 51 | PA2679 | 208964.PA2679 | 1 |
| 52 | PA3235 | 208964.PA3235 | 1 |
| 53 | PA3420 | 208964.PA3420 | 1 |
| 54 | PA3441 | 208964.PA3441 | 1 |
| 55 | PA3694 | 208964.PA3694 | 1 |
| 56 | PA3882 | 208964.PA3882 | 1 |
| 57 | PA5328 | 208964.PA5328 | 1 |
| 58 | *hutU* | 208964.PA5100 | 1 |
| 59 | *ospR* | 208964.PA2825 | 1 |
| 60 | *pitA* | 208964.PA4866 | 1 |
| 61 | *pslK* | 208964.PA2241 | 1 |
| 62 | *ureD* | 208964.PA4864 | 1 |

Rest 58 genes with node degree score ‘zero’ are not listed.

**Table S6. Top fourteen cytoHubba ranked genes from among the top-26 in Table S5**

| **No.** | **Gene ID** | **Gene Name** | **Number of methods ranking this protein among top 10** | **Names of 12 ranking methods of CytoHubba and rank score provided by them** | | | | | | | | | | | |
|---|---|---|---|---|---|---|---|---|---|---|---|---|---|---|---|
| | | | | **Degree** | **MNC** | **DMNC** | **MCC** | **Bottleneck** | **EcCentricity** | **Closeness** | **Radiality** | **Betweenness** | **Stress** | **CC** | **EPC** |
| 1 | PA2357 | *msuE, slfA* | 11 | 4 | 3 | 0.46346 | 7 | 6 | 0.10256 | 5.333333 | 1.27473 | 9 | 18 | - | 5.113 |
| 2 | PA3442 | *ssub1* | 10 | 4 | 4 | 0.47366 | 12 | 1 | - | 5.083333 | 1.18681 | 0.5 | 2 | - | 5.131 |
| 3 | PA0185 | *atsB* | 9 | 4 | 4 | 0.47366 | 12 | - | - | 5.083333 | 1.18681 | 0.5 | 2 | - | 5.124 |
| 4 | PA3938 | *tauA* | 8 | 4 | - | - | 7 | 2 | - | 5.333333 | 1.27473 | 9 | 18 | - | 5.101 |
| 5 | PA4597 | oprJ | 11 | 4 | 3 | 0.46346 | 7 | 2 | 0.19231 | 4 | 0.52885 | 6 | 6 | - | 3.575 |
| 6 | PA4599 | mexC | 11 | 3 | 3 | 0.46346 | 6 | 1 | - | 3.5 | 0.48077 | 0 | 0 | 1 | 3.436 |
| 7 | PA0630 | | 10 | 4 | 4 | - | 8 | 1 | 0.19231 | 4 | 0.52885 | 2 | 4 | - | 3.827 |
| 8 | PA0807 | ampDh3 | 10 | 4 | 4 | - | 8 | 2 | 0.19231 | 4 | 0.52885 | 2 | 4 | - | 3.835 |
| 9 | PA2090 | | 9 | 4 | 4 | 0.47366 | 12 | - | - | 5.083333 | 1.18681 | 0.5 | 2 | - | 5.094 |
| 10 | PA3287 | | 9 | 3 | | 0.46346 | 6 | 1 | - | - | - | 0 | 0 | 1 | 3.456 |
| 11 | PA4596 | | 9 | 3 | 3 | 0.46346 | 6 | 1 | - | 3.5 | 0.48077 | - | - | 1 | 3.456 |
| 12 | PA5445 | | 9 | 3 | 3 | 0.46346 | 6 | 1 | 0.15385 | - | - | 0 | 0 | 1 | - |
| 13 | PA2555 | | 7 | 3 | 3 | 0.46346 | 6 | 1 | 0.15385 | - | - | - | - | 1 | - |
| 14 | PA3383 | | 7 | - | - | - | - | 3 | 0.153846 | 5 | 1.27473 | 20.5 | 34 | - | 4.629 |

"-": This method did not rank the shown protein among top 10

MNC: Maximum Neighborhood Component; DMNC: Density of Maximum Neighborhood Component; MCC: Maximal Clique Centrality; CC: Clustering Co-efficient; EPC: Edge Percolated Component

**Table S7. List of DEG in Herboheal-exposed *S. aureus* satisfying the dual criteria of log fold-change ≥2 and FDR≤0.01**

| No. | Feature ID/ Gene | Coding for | log fold change | FDR | Up- or down-regulation |
|---|---|---|---|---|---|
| 1 | *sarT* | HTH-type transcriptional regulator SarT | 18.00 | 0.005 | ↑ |
| 2 | SAFDA_1030 | Alpha-hemolysin | 17.35 | 0 | ↓ |
| 3 | SAFDA_1326 | Hypothetical protein | 11.00 | 0.003 | ↓ |
| 4 | SAFDA_0523 | Hypothetical protein | 10.00 | 0.006 | ↓ |
| 5 | SAFDA_0033 | Hypothetical protein | 8.50 | 0.01 | ↑ |
| 6 | *hlgA* | Gamma-hemolysin component A precursor | 7.16 | 2.45E-14 | ↓ |
| 7 | SAFDA_1218 | Sensor histidine kinase | 6.91 | 8.32E-07 | ↓ |
| 8 | SAFDA_1829 | Truncated beta-hemolysin | 6.75 | 0.003 | ↓ |
| 9 | SAFDA_0271 | Pyrimidine nucleoside transporter (nupC) | 6.16 | 2.27E-06 | ↑ |
| 10 | SAFDA_1022 | Fibrinogen binding-related protein | 6.00 | 6.10E-05 | ↓ |
| 11 | SAFDA_1441 | Competence protein ComGA | 6.00 | 0.007 | ↑ |
| 12 | SAFDA_2337 | Hypothetical protein | 5.87 | 0 | ↓ |
| 13 | *hlgB* | Gamma-hemolysin component B precursor | 5.83 | 4.33E-15 | ↓ |
| 14 | SAFDA_0231 | Hypothetical protein | 5.50 | 0.01 | ↑ |
| 15 | SAFDA_1217 | ABC transporter permease | 5.44 | 0.0002 | ↓ |
| 16 | *acuA* | Acetoin utilization protein | 4.80 | 0.01 | ↓ |
| 17 | *icaR* | Intercellular Adhesin Locus Regulator | 4.80 | 0.01 | ↓ |
| 18 | *ureA* | Urea catabolic process | 4.80 | 0.01 | ↑ |
| 19 | SAFDA_0372 | Hypothetical protein | 4.67 | 0.001 | ↓ |
| 20 | SAFDA_1187 | Hypothetical protein | 4.60 | 0.007 | ↑ |
| 21 | *saeP* | | 4.26 | 1.77E-07 | ↓ |
| 22 | SAFDA_0127 | Hypothetical protein | 4.25 | 0.004 | ↑ |
| 23 | SAFDA_2543 | Hypothetical protein | 4.13 | 2.77E-06 | ↑ |
| 24 | *saeR* | two-component system, OmpR family, response regulator | 4.10 | 1.68E-07 | ↓ |
| 25 | SAFDA_0843 | HAD superfamily hydrolase | 4.00 | 0.0006 | ↓ |
| 26 | *glpQ* | Glycerophosphoryldiesterphosphodiesterase | 3.85 | 0.0009 | ↓ |
| 27 | *dapB* | 4-hydroxy-tetrahydrodipicolinate reductase | 3.71 | 0.01 | ↓ |
| 28 | *ureD* | urease accessory protein | 3.66 | 6.33E-05 | ↑ |
| 29 | SAFDA_1182 | Phage repressor | 3.64 | 0.004 | ↓ |

| No. | Feature ID/ Gene | Coding for | log fold change | FDR | Up- or down-regulation |
|---|---|---|---|---|---|
| 30 | *hlgC* | Gamma-hemolysin component C precu | 3.63 | 4.79E-08 | ↓ |
| 31 | *modC* | molybdenum transport protein | 3.57 | 1.41E-05 | ↑ |
| 32 | SAFDA_1138 | 50S ribosomal protein L7 | 3.54 | 0.002 | ↓ |
| 33 | *saeS* | Two-component system, OmpR family, sensor histidine kinase | 3.47 | 3.31E-10 | ↓ |
| 34 | *secG* | Preproteintranslocase subunit | 3.44 | 0.001 | ↓ |
| 35 | *splA* | Serine protease | 3.44 | 0.001 | ↓ |
| 36 | *sbi* | Immunoglobulin G-binding protein Sbi | 3.43 | 1.02E-09 | ↓ |
| 37 | SAFDA_0853 | Hypothetical protein | 3.38 | 0.003 | ↓ |
| 38 | SAFDA_0003 | S4 region YaaA family protein | 3.33 | 0.002 | ↓ |
| 39 | SAFDA_1229 | Hypothetical protein | 3.33 | 0.01 | ↓ |
| 40 | *trpA* | Tryptophan synthase alpha chain | 3.33 | 0.01 | ↑ |
| 41 | SAFDA_1219 | Two-component response regulator | 3.26 | 6.13E-06 | ↓ |
| 42 | SAFDA_0085 | Hypothetical protein | 3.22 | 0.001 | ↑ |
| 43 | SAFDA_0794 | Hypothetical protein | 3.22 | 0.001 | ↑ |
| 44 | SAFDA_0277 | Hypothetical protein | 3.18 | 0.002 | ↑ |
| 45 | SAFDA_1494 | HAD superfamily hydrolase | 3.14 | 0.005 | ↓ |
| 46 | SAFDA_1538 | Hypothetical protein | 3.13 | 0.004 | ↑ |
| 47 | SAFDA_0193 | Hypothetical protein | 3.05 | 0 | ↓ |
| 48 | SAFDA_0562 | Hydrolase | 2.93 | 0.007 | ↓ |
| 49 | SAFDA_1410 | Hypothetical protein | 2.93 | 2.62E-07 | ↓ |
| 50 | SAFDA_2187 | Phosphosugar-binding transcriptional regulator | 2.92 | 0.01 | ↑ |
| 51 | SAFDA_t0025 | tRNA-Cys | 2.88 | 0.007 | ↓ |
| 52 | *spsB* | signal peptidase I | 2.88 | 0.007 | ↓ |
| 53 | SAFDA_0228 | Choloylglycine hydrolase | 2.87 | 0.007 | ↑ |
| 54 | *glpP* | Glycerol uptake operon antiterminator regulatory protein | 2.85 | 0.01 | ↑ |
| 55 | *lukG* | leukocidin/hemolysin toxin family protein | 2.80 | 9.14E-06 | ↓ |
| 56 | SAFDA_2043 | Hypothetical protein | 2.78 | 6.55E-10 | ↓ |
| 57 | SAFDA_r0007 | 5S ribosomal RNA | 2.71 | 0 | ↑ |
| 58 | *coaE* | dephospho-CoA kinase | 2.71 | 0.0008 | ↑ |
| 59 | SAFDA_0232 | Hypothetical protein | 2.71 | 0.01 | ↑ |
| 60 | *pnp* | Polyribonucleotide nucleotidyltransferase | 2.71 | 4.39E-08 | ↑ |
| 61 | SAFDA_2297 | Hypothetical protein | 2.66 | 0.01 | ↑ |
| 62 | SAFDA_2405 | MmpL efflux pump | 2.66 | 9.41E-09 | ↑ |

| No. | Feature ID/ Gene | Coding for | log fold change | FDR | Up- or down-regulation |
|---|---|---|---|---|---|
| 63 | SAFDA_0565 | Alpha/beta fold family hydrolase | 2.66 | 6.40E-11 | ↑ |
| 64 | SAFDA_2223 | ABC transporter permease | 2.64 | 0.01 | ↑ |
| 65 | SAFDA_0423 | Orn Lys Arg decarboxylase family protein | 2.63 | 1.62E-08 | ↑ |
| 66 | SAFDA_2221 | Hypothetical protein | 2.63 | 0.008 | ↑ |
| 67 | *tagX* | glycosyltransferase | 2.58 | 0.002 | ↓ |
| 68 | *sraP* | Serine-rich adhesin for platelets | 2.55 | 7.48E-10 | ↑ |
| 69 | SAFDA_2160 | Transcription regulator | 2.55 | 0.0003 | ↑ |
| 70 | SAFDA_0932 | Hypothetical protein | 2.53 | 1.10E-05 | ↓ |
| 71 | SAFDA_1828 | Truncated cell surface protein map-w | 2.52 | 7.00E-07 | ↓ |
| 72 | *saeQ* | transmembrane protein | 2.47 | 0.001 | ↓ |
| 73 | *drm* | | 2.45 | 0 | ↑ |
| 74 | SAFDA_0392 | Cobalamin synthesis protein | 2.45 | 0.01 | ↑ |
| 75 | SAFDA_2310 | Amino acid transporter | 2.43 | 0.008 | ↑ |
| 76 | SAFDA_2453 | 2-dehydropantoate 2-reductase | 2.43 | 0.002 | ↑ |
| 77 | *sarA* | Transcriptional regulator SarA | 2.43 | 2.11E-07 | ↑ |
| 78 | SAFDA_1135 | Hypothetical protein | 2.43 | 0.003 | ↓ |
| 79 | *nreA* | | 2.41 | 0.005 | ↓ |
| 80 | *gcvH* | Glycine cleavage system H protein | 2.33 | 0.0004 | ↓ |
| 81 | *sdaAB* | L-serine dehydratase | 2.32 | 0.006 | ↓ |
| 82 | *recQ_1* | ATP-dependent DNA helicase RecQ | 2.32 | 4.68E-06 | ↑ |
| 83 | SAFDA_2273 | Polar amino acid ABC transporter ATPase | 2.31 | 0.003 | ↓ |
| 84 | *icaA* | intercellular adhesion (ica) locus | 2.30 | 0.009 | ↑ |
| 85 | SAFDA_0225 | Ribose transcriptional repressor RbsR | 2.28 | 0.001 | ↑ |
| 86 | SAFDA_2537 | Lipoprotein, putative | 2.26 | 0.01 | ↑ |
| 87 | *rpsO* | Small subunit ribosomal protein S15 | 2.26 | 9.03E-06 | ↓ |
| 88 | SAFDA_1759 | Sugar ABC transporter ATPase | 2.26 | 2.72E-07 | ↓ |
| 89 | SAFDA_0987 | Hypothetical protein | 2.25 | 0.009 | ↓ |
| 90 | SAFDA_2261 | Transcriptional regulator NirR | 2.25 | 0.009 | ↓ |
| 91 | SAFDA_0998 | Iron-regulated heme-iron binding protein | 2.25 | 0.001 | ↑ |
| 92 | SAFDA_1045 | HAD superfamily hydrolase | 2.25 | 0.009 | ↑ |
| 93 | *xerC* | Integrase/recombinase | 2.24 | 0.005 | ↓ |
| 94 | SAFDA_2230 | Glycosylglycerophosphatetransferase involvedin teichoic acid biosynthesis | 2.22 | 0.003 | ↑ |

| No. | Feature ID/ Gene | Coding for | log fold change | FDR | Up- or down-regulation |
|---|---|---|---|---|---|
| 95 | *pheT* | Phenylalanine--tRNA ligase beta subunit | 2.22 | 1.75E-07 | ↑ |
| 96 | SAFDA_2057 | Alcohol dehydrogenase | 2.21 | 5.04E-08 | ↑ |
| 97 | SAFDA_0091 | Major facilitator transporter | 2.21 | 6.48E-14 | ↑ |
| 98 | *tcaB* | teicoplanin-associated operon | 2.20 | 0.001 | ↑ |
| 99 | *dra* | | 2.19 | 5.62E-11 | ↑ |
| 100 | SAFDA_1716 | Hypothetical protein | 2.18 | 0.006 | ↓ |
| 101 | *Dps* | General stress protein 20U | 2.17 | 0 | ↓ |
| 102 | SAFDA_2462 | Hypothetical protein | 2.14 | 0.007 | ↑ |
| 103 | SAFDA_0189 | ABC transporter substrate-binding protein | 2.11 | 0.009 | ↑ |
| 104 | *Geh* | Glycerol ester hydrolase | 2.11 | 6.59E-06 | ↓ |
| 105 | *rpoE* | Probable DNA-directed RNA polymerase subunit delta | 2.10 | 0.001 | ↑ |
| 106 | SAFDA_1657 | Aesenical pump membrane protein | 2.10 | 0.005 | ↑ |
| 107 | *rpsB* | Small subunit ribosomal protein S2 | 2.09 | 3.60E-11 | ↓ |
| 108 | SAFDA_2315 | M42 glutamylaminopeptidase, cellulose | 2.04 | 4.16E-10 | ↑ |
| 109 | *ureC* | Urease subunit alpha | 2.03 | 0.0008 | ↑ |
| 110 | *mviM* | putative oxidoreductase | 2.02 | 0.01 | ↑ |
| 111 | SAFDA_1331 | Major facilitator superfamily permease | 2.02 | 0.01 | ↑ |
| 112 | *dapA* | 4-hydroxy-tetrahydrodipicolinate synthase | 2.01 | 0.001 | ↑ |
| 113 | SAFDA_2229 | L-lactate permease | 2.00 | 3.68E-09 | ↓ |

Genes are arranged in decreasing order of Fold Change.

**Table S8. Node degree score of up-down regulated genes mentioned in Table S7**

| No. | Gene symbol | Identifier | Node degree |
|---|---|---|---|
| 1 | *sbi* | 1280.SAXN108_2673 | 7 |
| 2 | *saeR* | 1280.SAXN108_0774 | 6 |
| 3 | *hlgB* | 1280.SAXN108_2677 | 5 |
| 4 | *hlgC* | 1280.SAXN108_2676 | 5 |
| 5 | *pheT* | 1280.SAXN108_1134 | 5 |
| 6 | *saeS* | 1280.SAXN108_0773 | 5 |
| 7 | *sarA* | 1280.SAXN108_0683 | 5 |
| 8 | *icaA* | 1280.SAXN108_2939 | 4 |
| 9 | *splA* | 1280.SAXN108_1846 | 4 |
| 10 | *lip2* | 1280.SAXN108_0305 | 3 |
| 11 | *pnp* | 1280.SAXN108_1278 | 3 |
| 12 | *rpsB* | 1280.SAXN108_1258 | 3 |
| 13 | *rpsO* | 1280.SAXN108_1277 | 3 |
| 14 | *icaR* | 1280.SAXN108_2938 | 2 |
| 15 | *ureA* | 1280.SAXN108_2536 | 2 |
| 16 | *ureC* | 1280.SAXN108_2538 | 2 |
| 17 | *ureD* | 1280.SAXN108_2542 | 2 |
| 18 | *coaE* | 1280.SAXN108_1714 | 1 |
| 19 | *dapA* | 1280.SAXN108_1411 | 1 |
| 20 | *dapB* | 1280.SAXN108_1412 | 1 |
| 21 | *sarT* | 1280.SAXN108_2745 | 1 |
| 22 | *tagX* | 1280.SAXN108_0708 | 1 |
| 23 | *trpA* | 1280.SAXN108_1389 | 1 |

Rest 5 genes with node degree score ‘zero’ are not listed.

### Table S9. Top twelve cytoHubba ranked genes from among the top-13 in Table S8

| No. | Gene ID | Gene Name | Number of methods ranking this protein among top 10 | Names of 12 ranking methods of CytoHubba and rank score provided by them | | | | | | | | | | | |
|---|---|---|---|---|---|---|---|---|---|---|---|---|---|---|---|
| | | | | Degree | MNC | DMNC | MCC | Bottleneck | EcCentricity | Closeness | Radiality | Betweenness | Stress | CC | EPC |
| 1 | SAXN108_0683 | *sarA* | 11 | 4 | 3 | 0.46346306 | 7 | 1 | 0.346154 | 6 | 2.076923 | 7.4 | 18 | - | 4.534 |
| 2 | SAXN108_2673 | *sbi* | 12 | 7 | 7 | 0.40246305 | 38 | 4 | 0.346154 | 7.5 | 2.336538 | 13.266667 | 28 | 0.52381 | 5.374 |
| 3 | SAXN108_1846 | *splA* | 11 | 4 | 4 | - | 8 | 1 | 0.346154 | 6 | 2.076923 | 2.8 | 8 | 0.666667 | 4.527 |
| 4 | SAXN108_0774 | *saeR* | 11 | 5 | 5 | 0.51861011 | 30 | - | 0.346154 | 6.5 | 2.163462 | 2.1333333 | 8 | 0.8 | 4.973 |
| 5 | SAXN108_0773 | *saeS* | 11 | 5 | 5 | 0.51861011 | 30 | - | 0.346154 | 6.5 | 2.163462 | 2.1333333 | 8 | 0.8 | 4.943 |
| 6 | SAXN108_2677 | *hlgB* | 11 | 5 | 5 | 0.51861011 | 30 | 1 | - | 6.333333 | 2.076923 | 1.3333333 | 4 | 0.8 | 4.876 |
| 7 | SAXN108_2676 | *hlgC* | 11 | 5 | 5 | 0.51861011 | 30 | 1 | - | 6.333333 | 2.076923 | 1.3333333 | 4 | 0.8 | 4 |
| 8 | SAXN108_1278 | *pnp* | 11 | 3 | 3 | 0.46346306 | 6 | 1 | 0.307692 | 3 | 0.512821 | 0 | 0 | 1 | - |
| 9 | SAXN108_0305 | *lip2* | 8 | 3 | - | - | - | 2 | 0.346154 | 5.5 | 1.990385 | 4.6 | 12 | - | 3.953 |
| 10 | SAXN108_1134 | *pheT* | 7 | 3 | 3 | 0.46346306 | 6 | 1 | 0.307692 | - | - | - | - | 1 | - |
| 11 | SAXN108_1277 | *rpsO* | 7 | - | 3 | 0.46346306 | 6 | 1 | 0.307692 | - | - | - | - | 1 | 2.315 |
| 12 | SAXN108_2939 | *icaA* | 6 | - | - | - | - | 1 | - | 4.666667 | 1.730769 | 1 | 2 | - | 3.092 |

"-": This method did not rank the shown protein among top 10

MNC: Maximum Neighborhood Component; DMNC: Density of Maximum Neighborhood Component; MCC: Maximal Clique Centrality; CC: Clustering Co-efficient; EPC: Edge Percolated Componen
